# Mechanical Flexibility Enables DNA Origami to Overcome Steric Confinement in Mucus

**DOI:** 10.64898/2026.03.21.713045

**Authors:** Matteo Tollemeto, Emily Tsang, Marie Karen Tracy Hong Lin, Luca Mannino, Katharina Ribbeck, Kurt V. Gothelf, Anja Boisen

**Author notes:** **Corresponding authors:** Matteo Tollemeto.

## Abstract

Size exclusion within biological hydrogels imposes a fundamental constraint on the design of nanocarriers, limiting the transport of cargo-loaded and structurally complex materials through mucus barriers. While surface passivation strategies are commonly used to improve compatibility, they do not address steric limitations imposed by the polymer network.

Here, we introduce mechanical flexibility as an independent materials design parameter to expand the functional transport window of nanocarriers in mucus. Using programmable DNA origami to decouple flexibility from size and surface chemistry, we show that increased structural compliance enhances transport under steric confinement by facilitating passage through confined network pores. When surface-driven aggregation dominates, passivation is required to restore transport, after which flexibility provides additional gains.

Together, these results establish mechanical flexibility as a general materials design strategy for improving transport under size-constrained conditions, with implications for nanocarrier engineering across biological barriers.

## 1. Introduction

Transport through mucus remains a central challenge in the design of effective mucosal drug delivery systems.^[1,2]^ While extensive work has examined how nanoparticle properties such as size,^[3,4]^ shape,^[5,6]^ and surface chemistry^[7,8]^ influence transport through mucus, two equally important factors remain comparatively underexplored: the mechanical flexibility of the nanostructure and the biological variability of the mucus barrier itself. Mucus is a heterogeneous viscoelastic network whose pore architecture and local mechanical constraints vary across length scales, suggesting that particle deformability could play an important role in navigating steric and adhesive barriers.^[9,10]^ However, flexibility has been difficult to investigate systematically because modifications that alter mechanical properties often simultaneously change particle geometry or surface chemistry, preventing the isolation of flexibility as an independent design parameter.

At the same time, most mechanistic studies evaluate particle transport in a single mucus source or in biosimilar mucus models,^[11,12]^ despite substantial differences in composition, microstructure, and biochemical complexity across anatomical regions and physiological states.^[13]^ Consequently, design principles derived from one mucus environment are often assumed to be broadly transferable, even though the dominant mechanisms governing mucoadhesion and transport may differ fundamentally between barriers.

To address these limitations, we employed a 14-helix bundle (14HB) DNA origami nanostructure as a programmable platform in which structural flexibility can be precisely tuned while maintaining nearly identical geometry and surface chemistry (**Fig. 1A, Fig. S1**).^[14,15]^ DNA origami enables nanoscale structural control through the folding of a long scaffold strand with hundreds of complementary staple strands.^[16,17]^ Building on previous work demonstrating that mechanical properties can be modulated through staple placement, flexibility was systematically increased by selectively removing staples within a central hinge region (**Fig. 1B**).^[15,18,19]^

**Figure 1.**
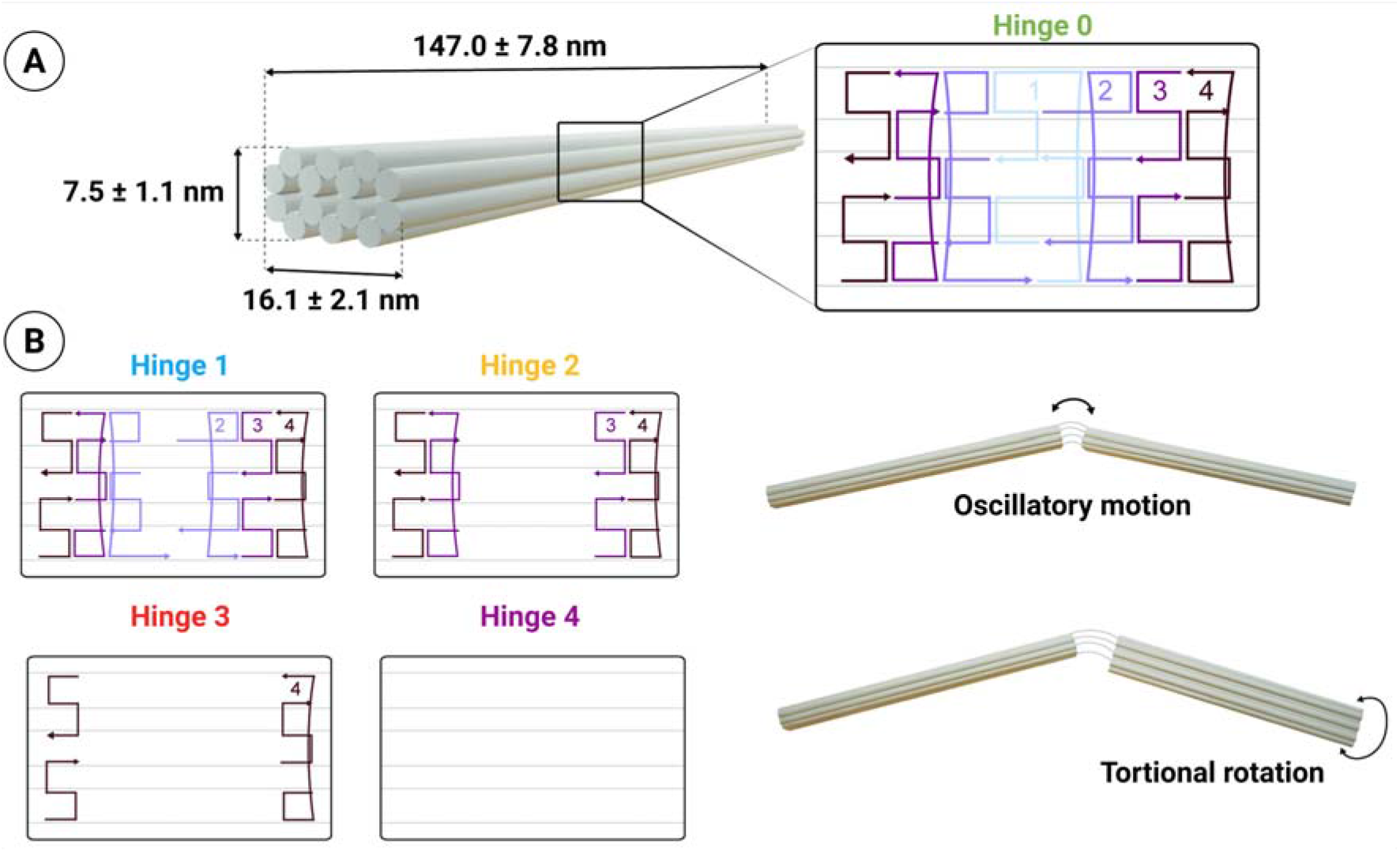
Modular DNA origami design and tunable hinge architecture. A) Schematic representation of a 14-helix bundle model system illustrating how structural modules define overall geometry and dimensions. B) The hinge element is generated by selectively omitting staple strands within the structural module, leaving unpaired scaffold single-stranded DNA at the cross-section. By varying the number of intact double-stranded segments and staple crossovers, the bending rigidity of the hinge can be systematically tuned. The resulting constructs exhibit distinct motion profiles corresponding to their programmed mechanical properties.

Using this platform, we investigated the influence of particle flexibility across three physiologically distinct mucus environments: fasted intestinal, fed intestinal, and stomach mucus. These samples differ in both their microstructural organization and biochemical composition, as confirmed by proteomic analysis. Comparing particle diffusion across these environments allowed us to distinguish between mechanically dominated and surface-interaction-driven transport limitations. When steric confinement within the polymer network represented the primary barrier, increased particle flexibility enhanced diffusion by facilitating passage through constricted pores. In contrast, in mucus environments where surface interactions promoted particle aggregation, surface passivation was required to mitigate mucoadhesion and restore mobility.

Finally, we demonstrate that prior exposure to ex vivo intestinal fluid significantly alters subsequent particle diffusion in mucus, indicating that interactions occurring in the fluid phase can strongly influence downstream transport behavior within the mucus layer. These findings highlight the importance of designing particles that are not only capable of penetrating mucus but also resistant to interactions with intestinal fluid components that may precondition their surface properties.

Together, this work demonstrates that optimal nanocarrier design requires identifying the dominant mechanisms governing transport in a given mucus environment. Whether steric obstruction, interfacial interactions, or aggregation limits mobility directly determines which design strategy, such as structural flexibility, surface passivation, or a combination of both, will be most effective for enhancing transport across mucosal barriers.^[20]^

## 2. Results and Discussion

### 2.1. Design, Structural Flexibility, and Diffusion Behavior of DNA Origami

Unlike parameters such as size, shape, or surface functionalization, structural flexibility cannot typically be tuned without altering the underlying architecture of a nanostructure.^[21]^ Consequently, modifications intended to change mechanical properties often introduce unintended variations in geometry or dimensions, complicating the systematic investigation of flexibility as an independent variable. DNA origami provides a unique solution to this challenge, enabling the construction of nanostructures with precisely defined shapes and dimensions while allowing mechanical properties to be tuned independently through selective staple placement.^[15,17]^ This programmability makes it possible to modulate structural flexibility without introducing confounding changes in size or surface chemistry, providing an ideal platform to study how mechanical properties alone influence interactions with complex biological environments.

To investigate this, we assembled a rod-shaped 14HB using the M13mp18 scaffold and systematically tuned its mechanical properties by selectively removing staples from the central “hinge” region. Progressive staple removal increased oscillatory bending, as supported by CanDo simulations (**Fig. S2**).^[22,23]^ This strategy generated five variants with progressively increasing flexibility, termed Hinge 0 to Hinge 4. Hinge 0 contained the complete staple set, whereas Hinge 4 incorporated the fewest staples and therefore exhibited the highest flexibility. Agarose gel electrophoresis showed no detectable differences in band mobility across the variants, indicating that staple removal did not significantly alter the overall structural dimensions (**Fig. S3**).

Transmission electron microscopy confirmed successful assembly of all designs and revealed a gradual increase in bending amplitude across the series (**Fig. 2A, Fig. S4-8**). Progressive staple removal in the hinge region resulted in a modest reduction in effective rod length (147.0 ± 7.8, 150.0 ± 4.6, 151.4 ± 4.7, 144.1 ± 3.6, and 136.8 ± 7.3 nm for H0–H4, respectively), determined by ImageJ analysis of more than 50 randomly selected structures. However, these differences were not statistically significant, indicating that flexibility remained the primary parameter systematically varied. Importantly, all variants retained their overall rod-like geometry, demonstrating that structural integrity was preserved despite staple removal.

**Figure 2.**
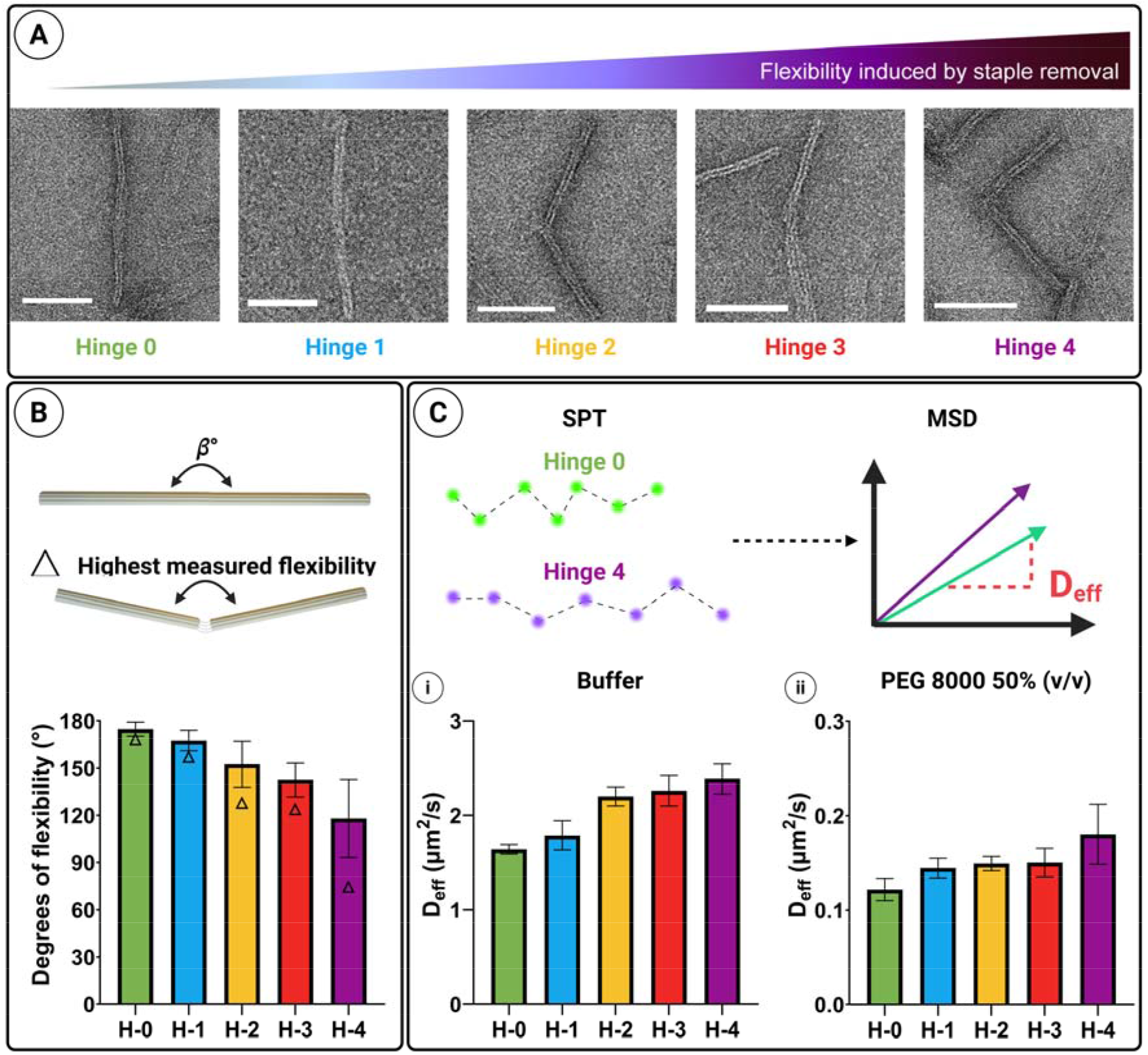
Programmable tuning of hinge flexibility and its impact on particle diffusion. A) Representative TEM micrographs of DNA origami constructs with increasing flexibility (Hinge 0–4), achieved by progressive staple removal. (Scale bars 100 nm). B) Quantification of hinge flexibility based on angular measurements (θ°). Top: schematic definition of the bending angle and illustration of the construct exhibiting the highest measured flexibility. Bottom: statistical analysis of bending angles for H-0 to H-4, demonstrating a systematic increase in structural compliance with hinge modification. C) Single-particle tracking (SPT) analysis of hinge constructs. Quantified D_eff_ values in (i) buffer and (ii) 50% (v/v) PEG 8000, showing flexibility-dependent differences in transport behavior (Table S1). (N > 30)

As expected, Hinge 0 exhibited a near-linear conformation with bending angles close to 180° (**Fig. 2B**). Increasing hinge flexibility from Hinge 1 to Hinge 4 produced a progressive decrease in bending angle, with Hinge 4 displaying a mean angle of approximately 120° and individual conformations as low as 75°. Notably, Hinge 3 and Hinge 4 also exhibited pronounced torsional motion, consistent with increased rotational freedom accompanying enhanced bending flexibility. Bending angles differed significantly between most variants (p < 0.05), except between Hinge 0 and Hinge 1 and between Hinge 2 and Hinge 3, confirming controlled and quantifiable modulation of mechanical flexibility without compromising structural assembly.

To determine how flexibility influences particle mobility independent of biological interactions, diffusion coefficients were measured for 100 individual particles in both buffer and a viscous inert medium (polyethylene glycol, PEG) that mimics the viscosity of mucus while eliminating specific interactions (**Fig. 2C**). In both environments, increased flexibility resulted in higher diffusion coefficients, consistent with the idea that bending and shape fluctuations reduce effective hydrodynamic drag and facilitate translational motion.^[24]^ Comparing the most rigid and most flexible designs, diffusivity increased by approximately 45% in buffer (H0: 1.64 ± 0.05 *µ*m^2^/s; H4: 2.39 ± 0.16 *µ*m^2^/s) and by approximately 48% in PEG (H0: 0.122 ± 0.012 *µ*m^2^/s; H4: 0.180 ± 0.032 *µ*m^2^/s). While all variants showed significant differences in buffer, no significant differences were observed between Hinge 1 and Hinge 2 or between Hinge 2 and Hinge 3 in PEG, consistent with the similar bending angles observed by TEM (**Table S1**). Based on these results, Hinge 0, Hinge 2, and Hinge 4 were selected for subsequent experiments to represent a broad range of mechanical flexibility and diffusion behavior in mucus.

### 2.2. Probing Mucus: Diffusion Dynamics and Biophysical Interactions

To examine the influence of structural flexibility on mucosal transport, particle diffusion was investigated in three ex vivo porcine mucus sources: fasted intestinal, fed intestinal, and fasted stomach mucus. These environments were selected to represent physiologically distinct mucus barriers that differ in composition, microstructure, and biochemical complexity.^[13]^ ATTO-647N–labeled 14HB DNA origami rods were tracked using an Oxford Nanoimager (ONI) microscope. For each mucus type, nanostructures were analyzed both with and without bovine serum albumin (BSA) conjugation to evaluate how structural flexibility and surface chemistry jointly influence particle transport. This experimental design enabled direct comparison of design-dependent transport behavior across biologically distinct mucus environments.

We first analyzed particle diffusion in fasted intestinal mucus (**Fig. 3A**). The absolute diffusion coefficients measured here were lower than those reported in our previous studies.^[25,26]^ This difference likely reflects variations in the biochemical composition and microstructural organization of the mucus samples used, highlighting the inherent biological variability of mucosal barriers. Despite these differences in absolute values, the overall trend remained consistent: more flexible DNA origami rods exhibited higher mobility and deeper penetration compared to their more rigid counterparts (**Fig. 3A i**). This observation supports the hypothesis that particle deformability facilitates transport through the sterically heterogeneous mucus mesh.

**Figure 3.**
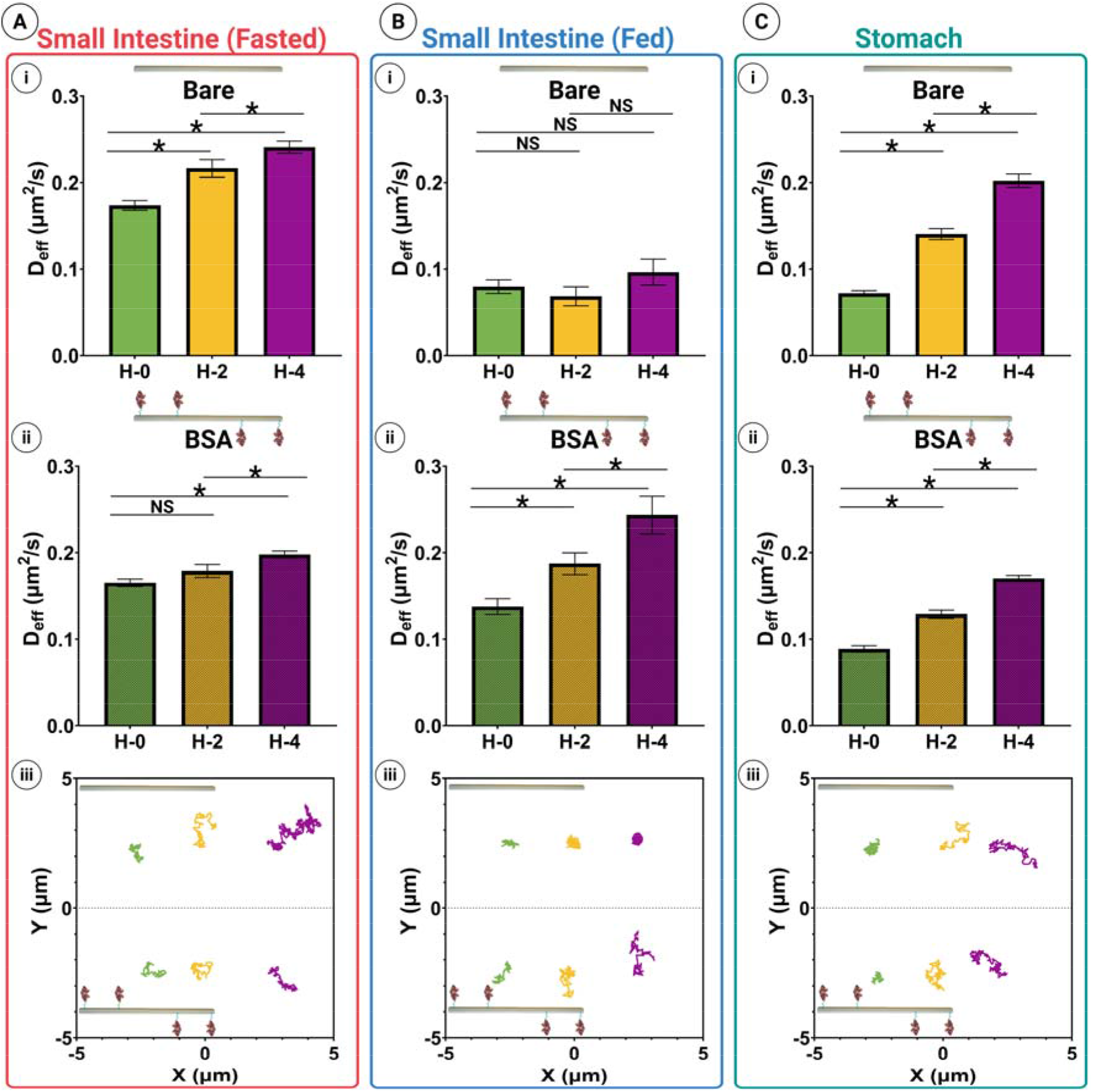
Flexibility effect on diffusion of DNA origami in mucus. Diffusion coefficients of nanoparticles in gastrointestinal mucus extracted from mean squared displacement (MSD) analysis. A) Intestinal mucus under fasted conditions: (i) particles of varying flexibility, (ii) BSA-coated particles, (iii) representative single-particle tracks. B) Intestinal mucus under fed conditions: (i) particles of varying flexibility, (ii) BSA-coated particles, (iii) single-particle tracks. C) Stomach mucus: (i) particles of varying flexibility, (ii) BSA-coated particles, (iii) single-particle tracks. Data are shown as the mean ± SEM (n = 100 particles). p < 0.05, NS: not significant.

Introducing protein conjugation produced a clear and reproducible effect. BSA passivation reduced the diffusion of all particle variants (**Fig. 3A ii**), consistent with previous observations that surface functionalization can increase hydrodynamic size and introduce additional steric interactions at the particle–mucus interface.^[26]^ Notably, while BSA conjugation reduced overall mobility, it did not eliminate the influence of mechanical flexibility. Flexible rods consistently retained a measurable diffusion advantage over rigid designs. These results indicate that although conjugation of protein cargo or functional motifs may inherently reduce particle mobility, structural flexibility remains an effective strategy for enhancing mucosal transport. Representative particle trajectories further illustrate the combined influence of flexibility and surface chemistry on transport behavior (**Fig. 3A iii**).

To assess how mucus composition influences these trends, the same experiments were performed in fed intestinal mucus (**Fig. 3B**) and stomach mucus (**Fig. 3C**). Compared to fasted intestinal mucus, an overall decrease in diffusion coefficients was observed in these environments (**Fig. 3B i**), suggesting stronger mucoadhesive interactions in both the fed intestinal and stomach mucus. Nevertheless, increased particle flexibility generally remained associated with higher diffusion, indicating that structural adaptability can partially compensate for stronger adhesive interactions within more complex mucus barriers.

### 2.3. Mucus Characterization: Linking Structure, Mechanics, and Protein Composition

To determine whether differences in particle transport could be attributed to variations in mucus properties, each sample was first characterized in terms of microstructure, mechanical properties, and biochemical composition. Cryogenic scanning electron microscopy (cryo-SEM) was used to visualize microstructural features (**Figure 4A**). Both intestinal mucus samples, regardless of feeding state, displayed similar macro- and microstructural organization, with comparable pore sizes and shapes. At higher magnification, fine network features were visible in both samples, although these structures appeared more pronounced in the fed intestinal mucus. In contrast, stomach mucus exhibited a markedly different architecture characterized by predominantly circular pores, whereas intestinal mucus contained more elongated, oval-shaped structures.

**Figure 4.**
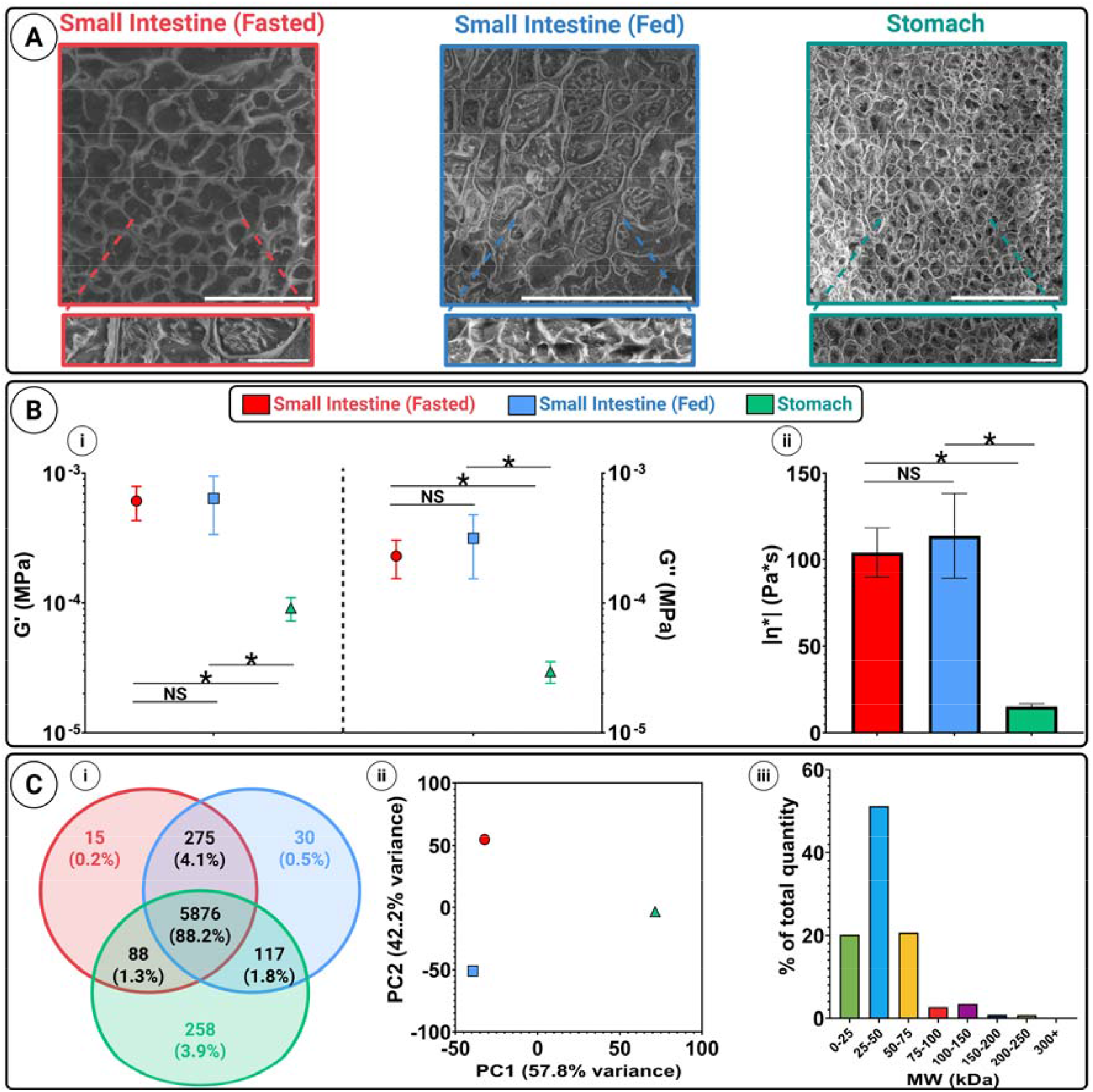
Structural profiling of porcine gastrointestinal mucus. A) Representative SEM micrographs of mucus samples collected from the small intestine under fasted and fed conditions, and from the stomach (Scale bars: 25∟*µ*m (top row), 5∟*µ*m (bottom row). B) Rheological properties of mucus from the small intestine (fasted and fed) and stomach. (i) Storage modulus (G′) and loss modulus (G″) on a logarithmic scale. (ii) Complex viscosity (|η*|). Data are presented as mean ± SD; *p < 0.05, NS: not significant. C) Proteomic analysis of mucus from the small intestine under fasted and fed conditions, and from the stomach. (i) total unique protein content, showing that most proteins are shared across samples; (ii) principal component analysis for the different mucus samples, highlighting sample-specific differences; (iii) molecular weight distribution of proteins uniquely identified in the fed intestinal sample, shown with abundance-weighted representation.

To further characterize material properties, rheological measurements were performed to determine the elastic modulus (G′), viscous modulus (G″), and complex viscosity (|η*|) of each mucus sample (**Fig. 4B**). Across all samples, the storage modulus (G′) remained lower than the loss modulus (G″), indicating that the measurements were performed within the linear viscoelastic regime and that no gelation occurred.^[13,27]^ Complex viscosity (|η*|), calculated from these moduli, provides an integrated measure of resistance to deformation and enables direct comparison between mucus samples. Both the elastic and viscous moduli were similar for fasted and fed intestinal mucus, although the fed samples exhibited larger standard deviations, suggesting greater heterogeneity likely arising from additional components such as digestive enzymes, bile salts, or dietary residues.^[28]^ In contrast, stomach mucus exhibited significantly higher moduli (p < 0.05) than both intestinal samples, indicating a mechanically stiffer viscoelastic network. Despite these differences, the measured complex viscosities fall within the range previously reported for ex vivo porcine intestinal mucus, confirming that the samples represent physiologically relevant conditions.^[11]^

To determine whether these structural and mechanical differences were associated with variations in biochemical composition, proteomic analysis was performed (**Fig. 4C**). Approximately 6300 proteins were identified in each sample, of which 5876 (88.2%) were shared across all mucus sources. Fasted intestinal mucus contained 15 unique proteins (0.2%), fed intestinal mucus contained 30 (0.5%), and stomach mucus contained 258 (3.9%). The large compositional overlap between fasted and fed intestinal mucus is consistent with their similar microstructure and rheological properties. In contrast, the greater number of unique proteins in stomach mucus correlates with its distinct structural organization. Principal component analysis further supported this interpretation: fasted and fed intestinal samples clustered closely along the first principal component, whereas stomach mucus separated clearly along this axis. Proteins contributing most strongly to principal components one and two are listed in **Table S2**.

Given the broadly similar overall protein molecular weight distributions across all samples, differences in structural organization are likely driven primarily by the subset of unique proteins. Although variations in the relative abundance of shared proteins could also influence network properties, weighted analyses of molecular weight distributions revealed only subtle differences between samples (**Figure S9**), suggesting that large abundance differences of overlapping proteins are unlikely to account for the observed structural variations. In fed intestinal mucus, the unique proteins were predominantly within the 25–50 kDa range, whereas unique proteins in fasted intestinal and stomach mucus were more broadly distributed across the full molecular weight range (0–300 kDa). The relatively small number of unique proteins in the intestinal samples is consistent with the comparable pore structures observed by cryo-SEM and their similar rheological properties. Despite these similarities, particle diffusion was slower in fed intestinal mucus, an effect that was largely mitigated by surface passivation. This observation suggests that the additional lower molecular weight proteins and associated biomolecules present in fed mucus may promote particle–mucus interactions without substantially altering the bulk polymer network. Under these conditions, transport limitation appears to be dominated primarily by mucoadhesive interactions.

In contrast, diffusion in stomach mucus improved only modestly following surface passivation, indicating that structural constraints play a more prominent role in limiting transport in this environment. The predominance of circular pores may impose stronger geometric restrictions on anisotropic particles. If this pore geometry persists at the nanoscale, which cannot be resolved by cryo-SEM, rod-shaped particles would be expected to align more readily within elongated pores, facilitating transport through the intestinal network. Circular pores, however, provide no preferred orientation, requiring particles to reorient during transport and thereby increasing the likelihood of transient confinement within the mucus mesh.

To better understand the mechanisms underlying the observed diffusion patterns, fluorescence imaging of individual particles was analyzed using single-particle tracking (SPT) (**Fig. 5**). Conventional bulk diffusion measurements could suggest that similar diffusion coefficients correspond to similar particle behavior in mucus.^[20]^ Under such an assumption, one might conclude that particles experience comparable mucoadhesive interactions across different mucus environments. However, SPT provides a more detailed view of particle dynamics, enabling direct visualization of particle–mucus interactions and identification of the dominant transport-limiting mechanisms.^[20]^

**Figure 5.**
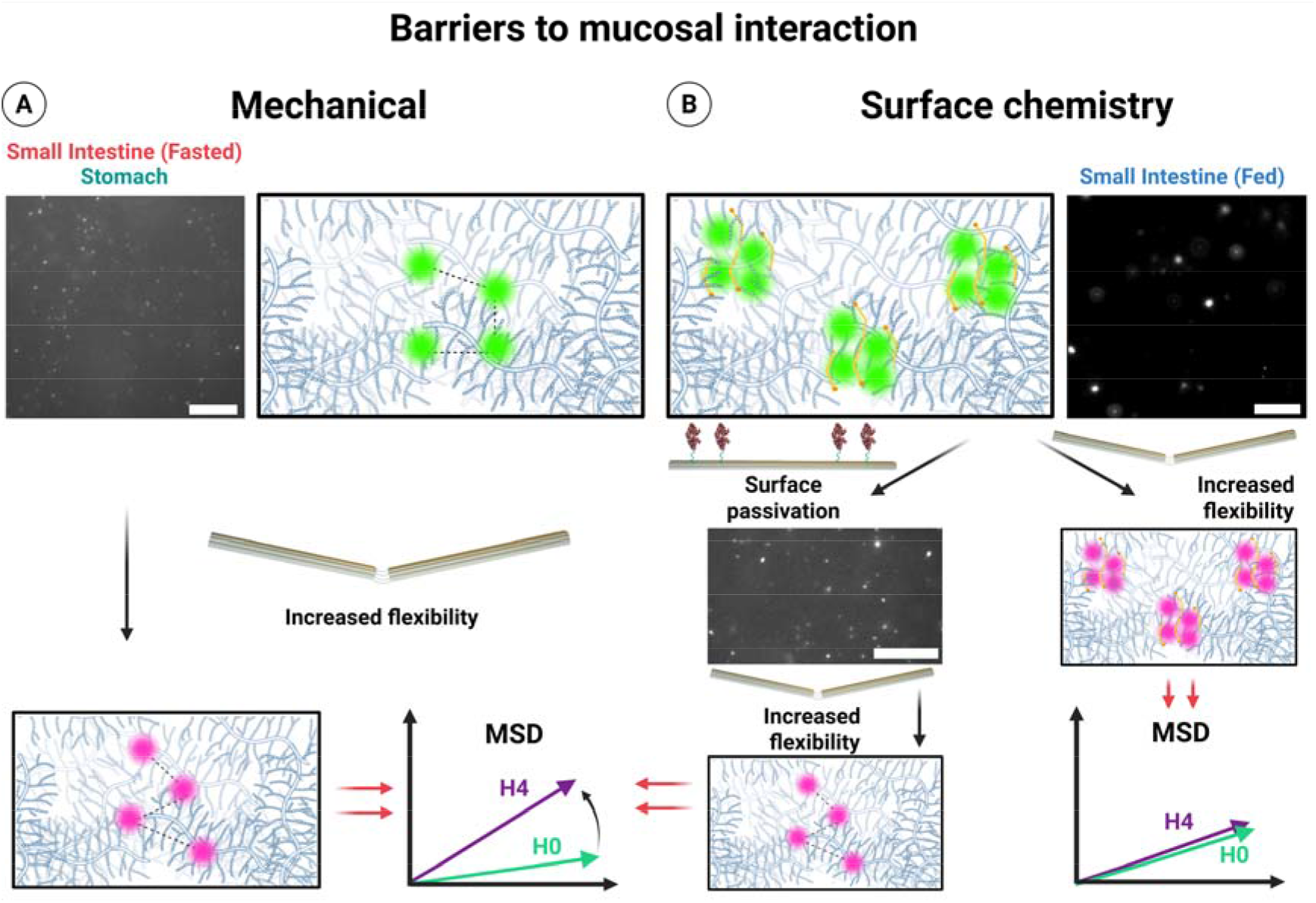
Schematic illustration of barriers to mucosal interaction and the effects of particle flexibility and surface chemistry. A) Mechanical barrier: Particles in fasted small intestine and stomach mucus experience steric hindrance. Increased particle flexibility allows deeper penetration into the mucin network. Corresponding mean squared displacement (MSD) plots illustrate enhanced diffusion for flexible particles (H4) compared to rigid particles (H0). B) Surface chemistry barrier: Particles in fed small intestine mucus experience aggregation. Increasing particle flexibility alone preserves aggregation, resulting in no significant change in diffusion MSD. Surface passivation reduces adhesive interactions with mucins, while increased flexibility further enhances mucus penetration (Scale bars: 10 μm).

Distinct particle behaviors were observed across mucus sources. In stomach and fasted intestinal mucus, particles predominantly appeared as individual entities with limited aggregation (**Fig. 5A**) In contrast, fed intestinal mucus exhibited pronounced particle clustering and aggregation (**Fig. 5B**). Despite comparable diffusion coefficients in some cases, these differences indicate that the underlying mechanisms governing transport are not equivalent. In stomach mucus, diffusion appears primarily limited by structural constraints such as steric hindrance within the polymer network. In fed intestinal mucus, however, interfacial interactions, including adsorption and electrostatic or hydrophobic interactions, promote particle aggregation and thereby reduce effective mobility.

These mechanistic differences have important implications for nanocarrier design. In environments where steric confinement dominates, such as stomach and fasted intestinal mucus, increasing particle flexibility enhances transport by facilitating passage through constricted network pores. In contrast, in fed intestinal mucus, where aggregation and surface interactions dominate, modifying particle flexibility alone has limited impact on diffusion. Instead, surface passivation becomes essential to prevent aggregation and restore particle mobility. Consistent with this interpretation, BSA coating significantly increased diffusion compared to unmodified particles. Moreover, once aggregation was suppressed, introducing structural flexibility further enhanced diffusion, demonstrating that flexibility becomes beneficial only after interfacial interactions are mitigated.

Representative SPT images further illustrate this effect, showing a substantial reduction in particle aggregates following BSA coating. Collectively, these observations demonstrate that particle transport through mucus is governed by a balance of steric, mechanical, and interfacial mechanisms. Identifying which of these mechanisms dominates in a given mucus environment provides a rational basis for selecting appropriate nanocarrier design strategies.

### 2.4. Sequential incubation in intestinal fluid and mucus to probe particle interactions

Finally, we investigated particle behavior under sequential exposure to ex vivo intestinal fluid followed by ex vivo mucus (**Fig. 6A**), a more physiologically relevant scenario that mimics the conditions encountered during oral delivery. Particle diffusion was first measured in intestinal fluid, where a noticeable reduction in mobility was observed compared to diffusion in aqueous buffer (**Fig. 6B**). This decrease indicates interactions with components present in the intestinal fluid that slow particle motion. Such interactions are important to consider, as intestinal fluid contains a complex mixture of enzymes, bile salts, and other macromolecules that can influence both drug solubility and particle–mucus interactions, ultimately affecting mucoadhesion and transport.^[29–31]^

**Figure 6.**
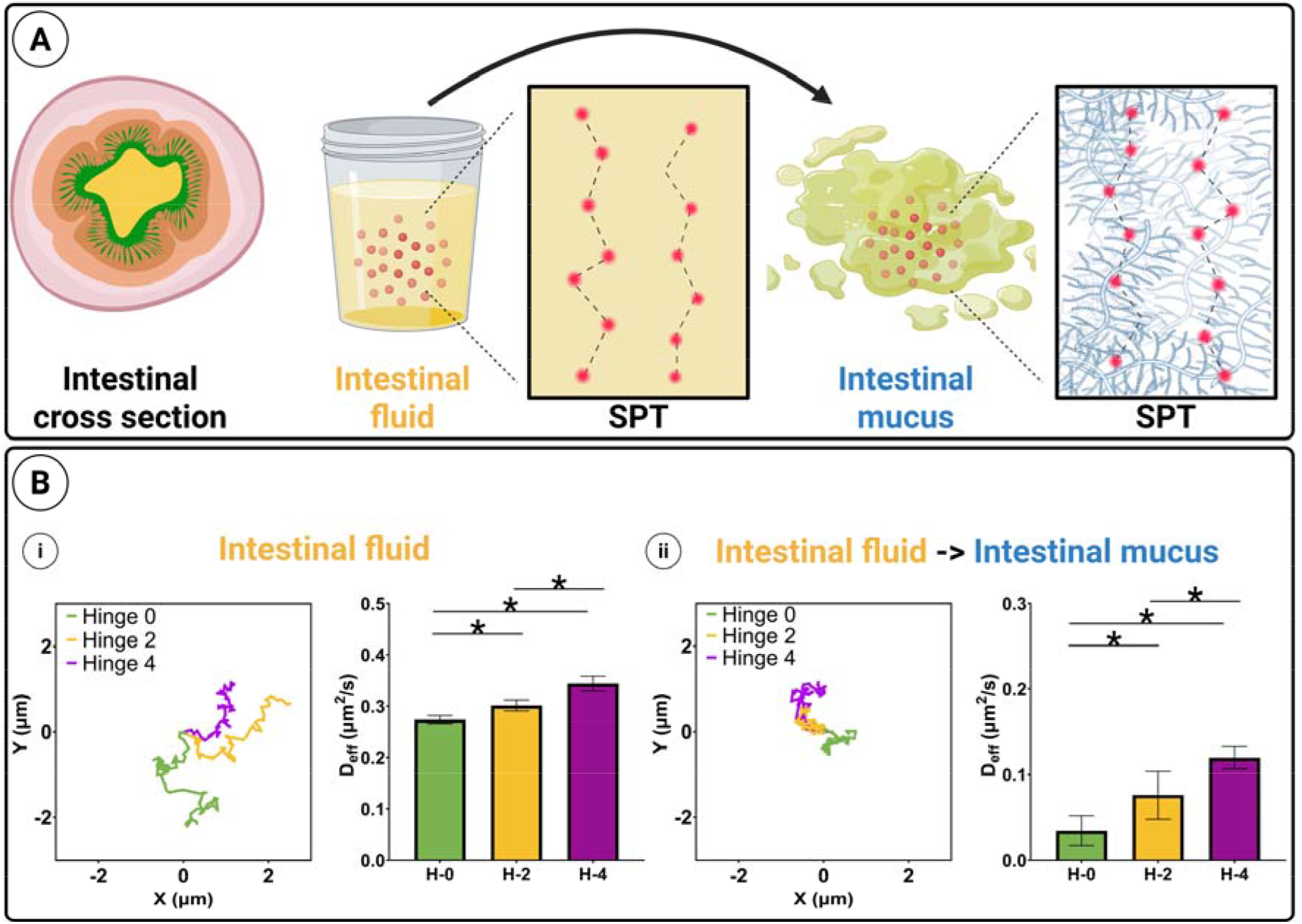
Sequential incubation. (A) Illustrative schematic of sequential incubation of particles first in intestinal fluid and then in mucus. (B) Representative particle tracks and Deff in intestinal mucus (i) and after sequential incubation in mucus (ii). Data are shown as the mean ± SEM (n = 100 particles). p < 0.05, NS: not significant.

Previous studies have shown that the DNA origami structures remain stable in intestinal mucus, ^[26]^ suggesting that structural degradation is not the primary factor responsible for the reduced diffusion observed here. Instead, interactions with fluid components likely alter particle surface properties or introduce transient binding events that hinder mobility. ^[32,33]^

Following incubation in intestinal fluid, particles were introduced into mucus to assess how prior fluid exposure influences subsequent transport. Compared to particles introduced directly from buffer, all variants exhibited further reduced diffusion coefficients after sequential incubation (**Fig. 6B ii**). Importantly, however, the relative effect of structural flexibility was preserved, with more flexible particles consistently showing higher mobility than rigid ones.

These results highlight that optimizing mucosal transport requires addressing interactions occurring both in intestinal fluid and within the mucus layer itself. While surface passivation strategies can reduce aggregation and adhesive interactions, mechanical flexibility remains an effective strategy for improving particle mobility once these interfacial effects are mitigated. Together, these findings demonstrate that rational design of mucopenetrative systems must account for sequential interactions with multiple biological barriers, including both intestinal fluid and mucus.

## 3. Conclusion

Classical strategies for designing mucopenetrating nanoparticles have largely relied on surface passivation to enhance diffusion,^[34,35]^ often without a complete understanding of the mechanisms governing particle–mucus interactions. While surface passivation can improve mucosal transport, our results demonstrate that optimal particle design requires first identifying the dominant factors limiting diffusion within a given mucus environment. Notably, widely used strategies such as PEGylation may compromise targeting and cellular uptake, highlighting the need for complementary design approaches.^[36]^

To address this challenge, we introduced particle flexibility as a design parameter that can be tuned independently of surface chemistry and overall geometry. Using DNA origami as a programmable platform, structural flexibility was precisely modulated while maintaining comparable shape, size, and surface properties. This enabled isolation of the mechanical contribution to mucosal transport. Increased flexibility markedly enhanced diffusion when transport was limited by steric confinement within the mucus network. In contrast, when diffusion was dominated by surface-driven interactions and particle aggregation, flexibility alone was insufficient; however, combining flexibility with surface passivation effectively mitigated these interactions and further improved penetration.

SPT was critical to resolve these mechanistic distinctions, as ensemble-averaged measurements would obscure aggregation and heterogeneity effects. Importantly, the transport advantages conferred by particle flexibility persisted under physiologically relevant conditions, where particles first interact with intestinal fluid prior to encountering the mucus layer. Sequential incubation studies revealed that fluid-mediated interactions can slow diffusion, but the relative benefits of flexibility remained, highlighting that effective mucopenetrative systems must address both fluid- and mucus-mediated transport limitations.

In summary, this work demonstrates that nanoparticle flexibility represents a previously underexplored yet powerful design parameter for mucosal drug delivery. By decoupling mechanical flexibility from particle geometry and surface chemistry using DNA origami, we show that structural compliance can significantly enhance transport in mucus environments where steric confinement dominates. In contrast, in systems governed by surface-mediated interactions and aggregation, flexibility alone is insufficient, and surface passivation must first mitigate adhesive interactions before mechanical advantages can be realized. These findings establish a mechanistic framework for rational nanocarrier design, in which particle properties are tailored to the dominant transport-limiting mechanisms within specific mucus environments. More broadly, this work highlights the importance of integrating structural mechanics with surface engineering to optimize nanoparticle transport across complex biological barriers.

## Materials and Methods

### Materials

All chemicals were used as received unless otherwise stated. The single-stranded M13p18 scaffold for 14-helix bundle was purchased from Tilibit, while staple strands, handle-extended staple strands, modified DNA oligonucleotides, and fluorescently modified DNA oligonucleotides, were obtained from Integrated DNA Technologies. All other chemicals and reagents, including: DBCO-PEG_4_-NHS ester, BSA, phosphate-buffered saline (PBS), glycogen, dimethyl sulfoxide (DMSO), NaCl, sodium tetraborate, DMF, MgCl_2_, CaCl_2_, HEPES, ethanol, NaOAc, EDTA, PEG 8000, RNase-free MQ were purchased from Sigma-Aldrich (St. Louis, MO). Solvents were HPLC grade and used as received. Ultrapure water was used throughout the experiments and obtained from Merck Millipore Q-Pod system (Merck Group, Burlington (MA), U.S.A) with a 0.22□μm Millipore Express 40 filter (18.2□MΩ).

### Data and statistical analysis

Data analysis was performed with GraphPad Prism (Version 9.4.1 (681) Insight Partners, Graphpad Holdings, LLC, New York City (NY), U.S.A.). Data are presented as average and standard deviation (SD). Statistical analysis was performed for statistically significant differences (*p < 0.05) with GraphPad Prism using a one-way analysis of variance (ANOVA) (three or more independent populations). Proteomics analyses, including data processing and statistical testing were performed using R (version 4.3.1). The figures were created with Biorender.com. Image analysis and processing was done using the software ImageJ (version 1.53t, National Institutes of Health, USA).

## Methods

### Design and assembly of DNA origami

The 14-helix bundle was designed using cadnano software, ^**[17]**^ following the same design as in our previous work,^**[26]**^ with selective staple removal introduced here to create a hinge.

Further details are outlined in the **Fig. S1**, respectively. The structures were assembled by using a single-stranded M13p18 scaffold at concentrations ranging from 3-40 nM, accompanied by a 10-fold molar excess of the staple strands. The sequences of all oligonucleotides used are provided in **Tables S3-6**. The folding buffer contained 20 mM MgCl_2_, 50 mM NaCl, and 1x TAE, and bundles were annealed by using the following method: 95 °C for 1 minute, 70 °C for 3 minutes, 70-40 °C ramp (1 °C/30 min) and 20 °C hold. After assembly the structures were purified using 100 kDa Amicon Ultra-0.5 mL centrifugal filters into a the same assembly buffer.

### Visualization of DNA origami structures

A 1% agarose gel containing 12.5 mM MgCl_2_ and 1x TBE was used to analyze the DNA origami structures. The gels were eluted at 65 V for 90 min and pre-stained with 1x SYBR Safe (Invitrogen). The samples were used for TEM.

### BSA-DNA conjugation reaction

To produce the maleimide-modified DNA strand, amino-modified DNA (10 nmol in 50 μL of MQ) was mixed with 4-(N-maleimidomethyl) cyclohexanecarboxylic acid N-hydroxysuccinimide ester (SMCC, 598 nmol in 50 μL of dry DMF) and 0.25 μL of TEA. The sample was then incubated overnight on a shaker at room temperature. After, the solution was then mixed with 250 μL 96% EtOH, 14 μL NaOAc (3 M, pH 5.2) and 1 μL glycogen (20 mg/ml). The mixture was flash frozen and centrifuged (14,000 RPM) at 4 °C for 45 min. Lastly, the pellet was resuspended in MQ and purified by RP-HPLC with Phenomenex Clarity 3u Oligo-RP 50 mm × 4.6 mm column running a gradient of acetonitrile in TEAA buffer (0.1 M, pH 7). The fractions containing product were collected and lyophilized before being dissolved in MQ and stored at -20 °C until required. A 10-fold molar excess of the maleimide-DNA strand was mixed with BSA in a buffer containing 100 mM sodium phosphate supplemented with 550 mM NaCl, pH 7.2 overnight at 37 °C, the total reaction volume was 200 μL. After, the mono-labeled BSA-DNA conjugate was purified using HPLC with a DNAPac™ PA-100 BioLC 4 × 250 nm column (Thermo Scientific) running a gradient of 25 mM TRIS supplemented with 1M NaCl, pH 8, followed by a gradient of 25 mM TRIS, pH 8. After isolation, the conjugates were concentrated using 30 kDa Amicon Ultra-0.5 mL centrifugal filters into a buffer of 1x PBS. The concentration was calculated using the extinction coefficient of 43,824 M^−1^cm^−1^ at 279 nm. The purified origami structures (Hinge 0, 2 and 4) with protein capture extensions, were mixed with 8x molar equivalences per binding site and left to incubate overnight at room temperature.

### TEM

The purified sample was deposited onto a glow-discharged carbon-coated grid (400 mesh, Ted Pella) for 1 min. The grid was then blotted, dipped on 8 μL of MQ before being blotted again. The grid was immediately treated twice with 2% uranyl formate (4 μL) with a 30 second hold before blotting the second treatment. The grids were left to dry for 2 min. Imaging was conducted with a Tecnai G2 Spirit TEM, operated at 120 kV.

### Mucus isolation

Intestines or stomach from healthy fasted (18–24 h) or fed gilts (40–60 kg, 3–4 months, Danish Landrace) were obtained after experimental surgery. Immediately after euthanization, the jejunum or stomach were isolated. Sections were opened by a latitude cut and porcine mucus was isolated by gently scraping the mucosal surface. Mucus was kept on ice at all times and stored at −20 °C until use. Procedures were according to the authorization by Danish Veterinary and Food Administration (license number: 2020-15-0201-00610).

### Mucus Cryo-Scanning Electron Microscope (Cryo-SEM)

Mucus was mounted for cryo-SEM on a sample holder attached to a transfer rod, rapidly frozen by plunging into slushed liquid nitrogen at −210 °C, and transferred to the preparation chamber stage at −180 °C (Quorum PP2000 Cryo Transfer System). The frozen sample was cleaved with a cold knife (facilitating an exposed surface in the fractured sample), sublimated at −80 °C for 15 min, and coated with Pt at a current of 4.5 mA for 30 s. The sample was then transferred under vacuum to the SEM stage at −160 °C in the Field Emission Scanning Electron Microscope (FEI Quanta 200 ESEM FEG) and imaged at 10 kV using an ETD detector.

### Rheological properties of mucus

Rheological characterization of the samples was performed with an ARES-G2 rheometer (TA Instruments, US) set to 1% controlled strain mode, thus helping to stay within the linear viscoelastic regime. Samples were previously incubated at 37 and measurements were performed with a parallel plate geometry of 8 mm at room temperature and in the frequency range 10-0.1 Hz.

## Proteomics

### MS sample preparation

50uL of each sample were lysed in 1:3 lysis buffer (6M GdCl, 10mM TCEP, 40mM CAA, 50mM HEPES pH8.5) then boiled at 95°C for 5 minutes, after which they were sonicated on high for 5x 60 seconds in a Bioruptor sonication water bath (Diagenode) at 4oC then disrupted using a TissueLyser (Qiagen), gradually increasing the frequency from 3□Hz to 30□Hz over 1 minute to ensure thorough homogenization. Lysed samples were centrifuged at 1500*g for 5 mins, and supernatants were transferred to clean LoBind Eppendorf tubes. Protein concentration was determined by BCA rapid gold (Thermo). A total of 20□*µ*g of the samples was taken and diluted with 100□mM TEAB at pH 8.5 to achieve a final guanidine hydrochloride concentration below 4□M. For protein binding preparation, Sera-Mag SpeedBeads A and B (Fisher Scientific) were mixed 1:1 then washed three times with 200□*µ*L MS-grade water. After the final wash, the excess liquid was removed, and the beads were resuspended in MS-grade water. The prepared beads were added to the samples at a 1:20 ratio, followed by the addition of 100% ethanol to bring the final ethanol concentration to at least 60%. The samples were mixed thoroughly by pipetting and incubated at 800□RPM for 15 minutes to facilitate protein binding to the beads. After incubation, the supernatant was removed, and the beads were washed three times with 200□*µ*L of 80% ethanol. Following the washing steps, the beads were resuspended in 75□*µ*L of digestion buffer containing 10□ng/*µ*L trypsin (MS-grade, Sigma) in 100□mM TEAB. The samples were then sonicated for 30 seconds in a water bath sonicator and incubated overnight at 37□°C with shaking at 1000□RPM. After digestion, the supernatants were transferred to clean Protein LoBind tubes (Eppendorf) and acidified 1:1 with 2% trifluoroacetic acid (TFA) then desalted using a SOLA*µ* SPE plate (HRP, Thermo). Solvent applications were processed by centrifugation at 1500 RPM between each step. The filters were first activated with 200 *µ*L of 100% methanol, followed by 200 *µ*L of 80% acetonitrile, 0.1% formic acid. The filters were equilibrated twice with 200 *µ*L of 1% TFA, 3% acetonitrile, and then the samples were loaded. After washing the tips twice with 200 *µ*L of 0.1% formic acid, the peptides were eluted into clean Eppendorf tubes using 40% acetonitrile, 0.1% formic acid.

The eluted peptides were concentrated using a Savant SpeedVac SPD120 (Thermo Scientific and reconstituted in 12 *µ*L of Solution A* (2% acetonitrile, 1% TFA) for analysis. The peptides were then centrifuged at 18,000 × g for 10 minutes to remove any residual particulate matter. The supernatant was carefully transferred, and peptide concentration was measured using a Nanodrop DeNovix DS-11 FX+ spectrophotometer.

### MS Analysis

500 ng of peptides were loaded onto an *µ*PAC trapping column (Thermo Scientific, 3003438) connected in line with a 50 cm *µ*PAC NEO column (Thermo Scientific, 7500467) using 100% solvent A (0.1% formic acid in water) at 750 bar on a Vanquish NEO UHPLC system (Thermo). The column oven was operated at 50 °C.

Peptides were eluted over a 68-minute gradient ranging from 4% to 45% solvent B (80% acetonitrile, 0.1% formic acid) at a flow rate of 250 nL/min. The Orbitrap Exploris 480 mass spectrometer (Thermo Fisher Scientific) was operated using a DIA method. Full MS spectra were acquired at a resolution of 120,000 with a normalized AGC target of 300% or maximum injection time set to auto, over a scan range of 400–1000 m/z.

MS2 spectra were acquired at a resolution of 30,000 with an AGC target of 800% or maximum injection time set to auto and a normalized collision energy of 27. Isolation windows were set to 12 m/z with window placement optimization enabled, and the MS2 scan range was set to 145–1450 m/z. MS performance was verified for consistency using the Pierce™ HeLa Protein Digest Standard (Thermo Scientific), and chromatography was monitored to ensure reproducibility.

### Database search

Raw files were analyzed using Spectronaut™ (version 17.4). Spectra were matched against the Sus scrofa database from UniProt (taxon ID 9823). Dynamic modifications were set to oxidation (M) and acetylation of protein N-termini. Cysteine carbamidomethylation was set as a static modification. All results were filtered to a 1% FDR, and protein quantification was performed at the MS2 level. Protein groups were inferred using IDPicker.

### Data analysis

Protein-group quantification tables were imported into R and missing/NaN entries were set to zero. Proteins were considered as detected when its corresponding quantity was > 0, and these detected sets were used to visualize overlap among samples using a three-set Venn diagram (ggvenn), where each region represents shared and sample-specific protein-group identifications. For exploratory multivariate analysis, principal component analysis was performed on the protein-by-sample quantity matrix after applying a log10(x+1) transform to accommodate zeros and compress the dynamic range typical of proteomics intensities; proteins with zero variance across samples were removed, and principal component analysis was computed using centered and unit-variance scaled variables (prcomp with center=TRUE, scale=TRUE). Molecular-weight distributions were generated by mapping representative UniProt accessions for each protein group to sequence-derived mass annotations retrieved via UniProt.ws for Sus scrofa (taxId 9823), converting mass from Da to kDa, binning proteins into predefined molecular-weight ranges, and summarizing distributions either as count-based percentages (each detected protein group contributes equally) or as abundance-weighted percentages (protein-group quantities summed within each molecular-weight bin and normalized within sample); molecular-weight summaries were visualized as bar plots to compare how identification frequency and total quantified signal were distributed across molecular-weight bins within and between sample-defined protein subsets.

### Single Particle Tracking

The origami structures were annealed, with a 10-fold excess of protein capture extensions, and a 10-fold excess of the six ATTO 647N-DNA capture extensions. Additionally, a 50-fold molar excess of the ATTO 647N-modified DNA strand was used. After annealing, the structures were purified from the excess staple strands and excess fluorophore strands using 100 kDa Amicon Ultra-0.5 mL centrifugal filters. The buffer used contained 20 mM MgCl_2_, 50 mM NaCl, and 1x TAE. The structures were then diluted to 10 nM and mixed with an 8-fold molar excess of the BSA-DNA conjugates and left to incubate for 1 hour at room temperature. DNA⍰origami samples were prepared at 10□nM in folding buffer and diluted 1:10 immediately before measurement to give a final concentration of 1□nM. Images were acquired in a Nanoimager® (ONI, Oxford) using the NimOS software with a 640 nm laser (190 mW). The sample was illuminated using a highly inclined and laminated optical sheet (HiLo) at an angle of 47°, at 15% laser power. Fluorescence was recorded using an ONI 100x, 1.49 NA oil immersion objective and passed through a quad-band pass dichroic filter. Images were acquired onto a 425×518-pixel region (pixel size 0.117 μm) for 10 s at 100 fps. For each experimental condition, a total of at least 2 different biological samples were analyzed. All measurements were taken approximately 2 minutes after incubation. The results were filtered using the NimOS software with the following settings: maximum frame gap = 5, minimum distance between frames = 0.800 *µ*m, exclusion radius = 1.200 *µ*m, a minimum number of steps = 50, a minimum diffusion coefficient of 0.01 μm^2^s^−1^. With these parameters, track steps were extracted, and analyzed using a Python-based code1 to obtain the trajectories of the DNA origami (N = 100) and calculate the mean-squared displacement (MSD) with the following equation (2):

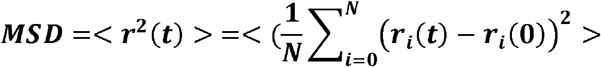

Then, the diffusion coefficient was obtained by fitting the MSD data to equation (3), where r = radius and t = sampling time and MSD(t) = 2dD, where D = diffusion coefficient and d = dimensionality (ONI measurements have dimension d = 2):

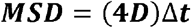

was used to fit the MSD curves.

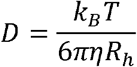

where *D* = diffusion coefficient, k_B_ = Boltzmann constant, *T* = temperature, η = mucus viscosity and Rh = hydrodynamic radius.^[37]^

To extract the α value, the MSD was plotted against lag time on a log–log scale, and a linear fit was applied; the slope of this fit corresponds to the anomalous diffusion exponent α.

## Acknowledgements

The authors would like to acknowledge the Danish National Research Foundation (DNRF122) and Villum Fonden (Grant No. 9301) for intelligent drug delivery and sensing using microcontainers and nanomechanics (IDUN), the Novo Nordisk Foundation (CEMBID: NNF17OC0028070, NNF17OC0026910 and NNF25OC0101151) and the European Research Council in Foldable, REconfigurable & Jagged devices for enhanced drug Absorption/seeding (FREJA), grant ref. No. 101054945.

The authors would like to acknowledge the Bioimaging Core Facility, Department of Biomedicine, Aarhus University, Denmark, for providing access to equipment and support with the ONI system. We also thank Prof. Thomas Thymann for assistance with tissue isolation, the DTU Proteomics Core Facility for running the proteomics samples, and Prof. Anne Ladegaard Skov and Dr. Liyun Yu for providing access to the rheometer.

