## Supporting Information for "Mechanical Flexibility Enables DNA Origami to Overcome Steric Confinement in Mucus"

**Corresponding authors:**

**Supplementary Figure 1.** Design schematic. (a) Illustration of the scaffold and staple routing for the flexible 14HB origami, as exported from Cadnano. (b) Color scheme indicating the different strands used in the assembly of the structure. (c) Front view of a simplified schematic showing the positions of strand extensions. (d) Simplified side view showing the position of protein capture extensions. (e) By removing more ‘hinge level’ staples, the hinge region (which is composed solely of the scaffold strand) becomes wider which increases its flexibility.

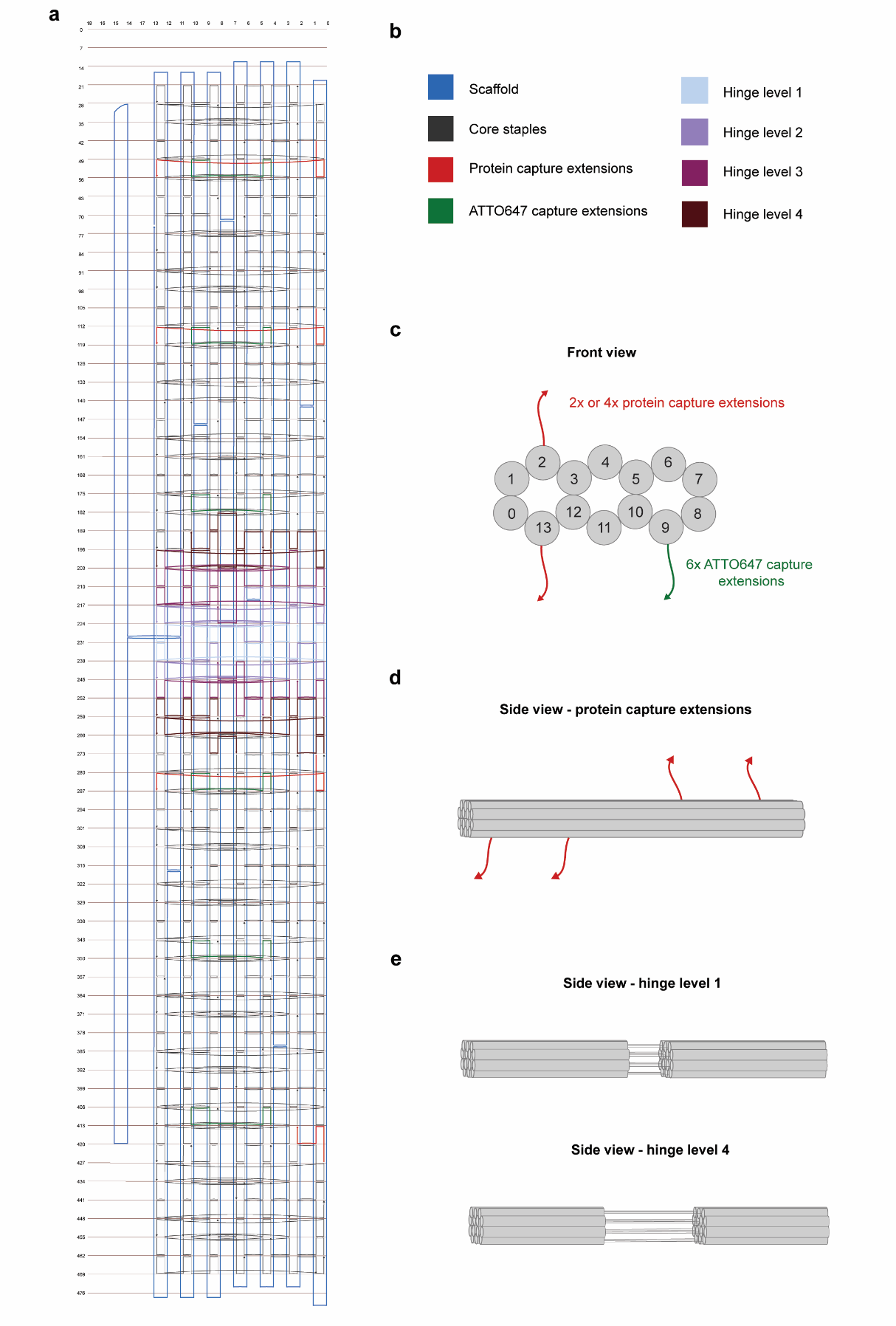

–

–

**Supplementary Figure 2.** Cando simulations demonstrate various flexibility around the hinge point for the different structures.

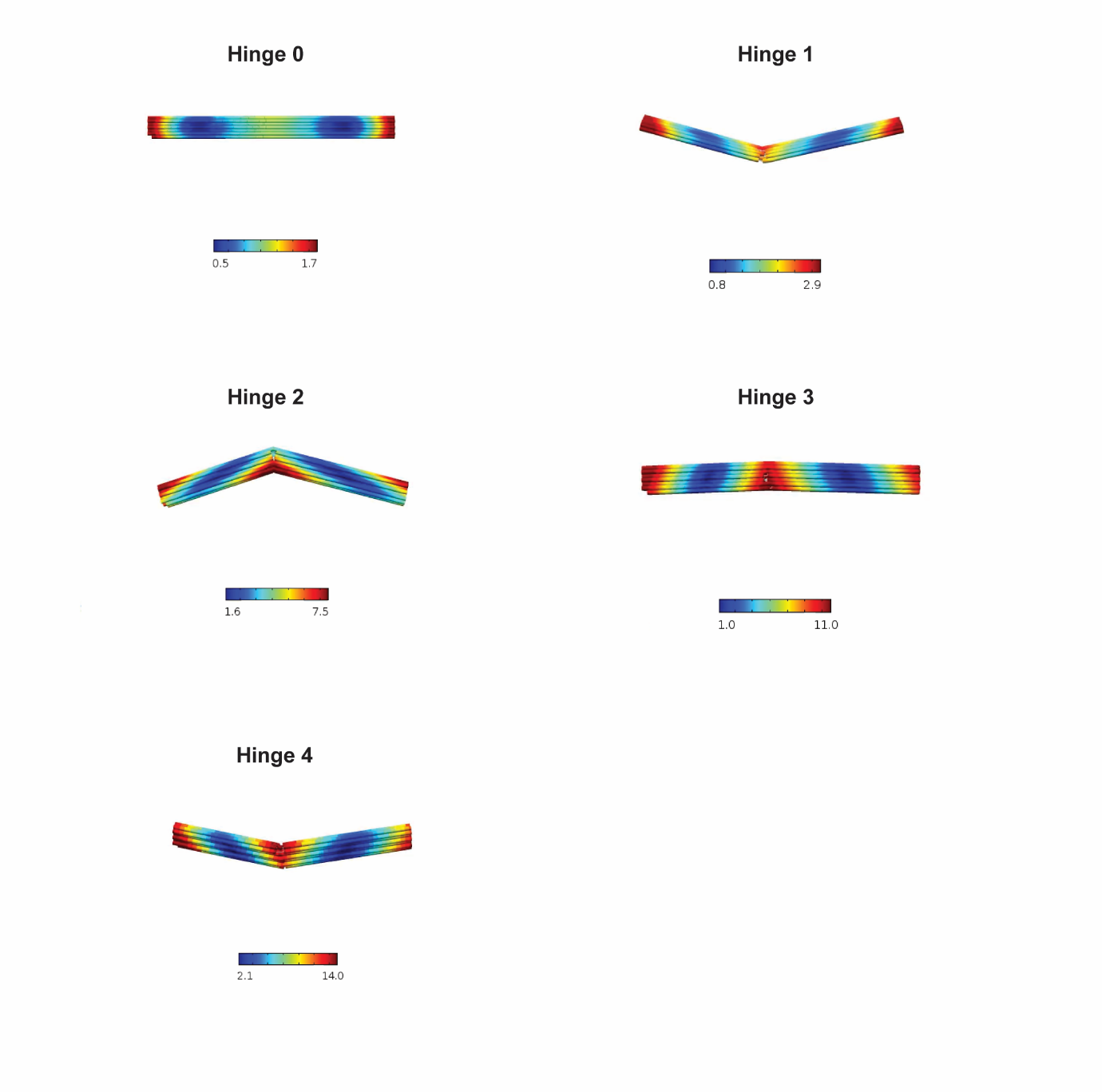

**Supplementary Figure 3.** Gel electrophoresis of the hinged structures. A 1% agarose gel (1x TBE, 5 mM MgCl_2_) was run at 65 V for 70 minutes, stained with SYBR Safe, and subsequently visualized. Increasing the ‘hinge level’ by removing additional staples produces progressively more flexible structures; however, only minimal differences in band mobility were observed.

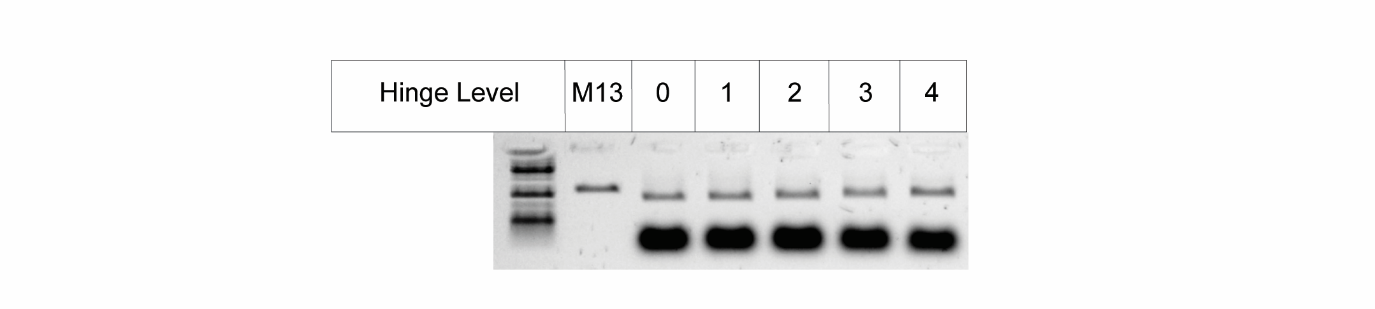

**Supplementary Figure 4.** Full TEM image of the 14HB with hinge 0. Before imaging, the structures were purified from the excess staples via Amicon filtration in a buffer containing 12.5 mM MgCl_2_, 50 mM NaCl, 1x TAE. Scale bar: 90 nm, magnification: 67,000x. A crop-out of this image is used in Figure 1b.

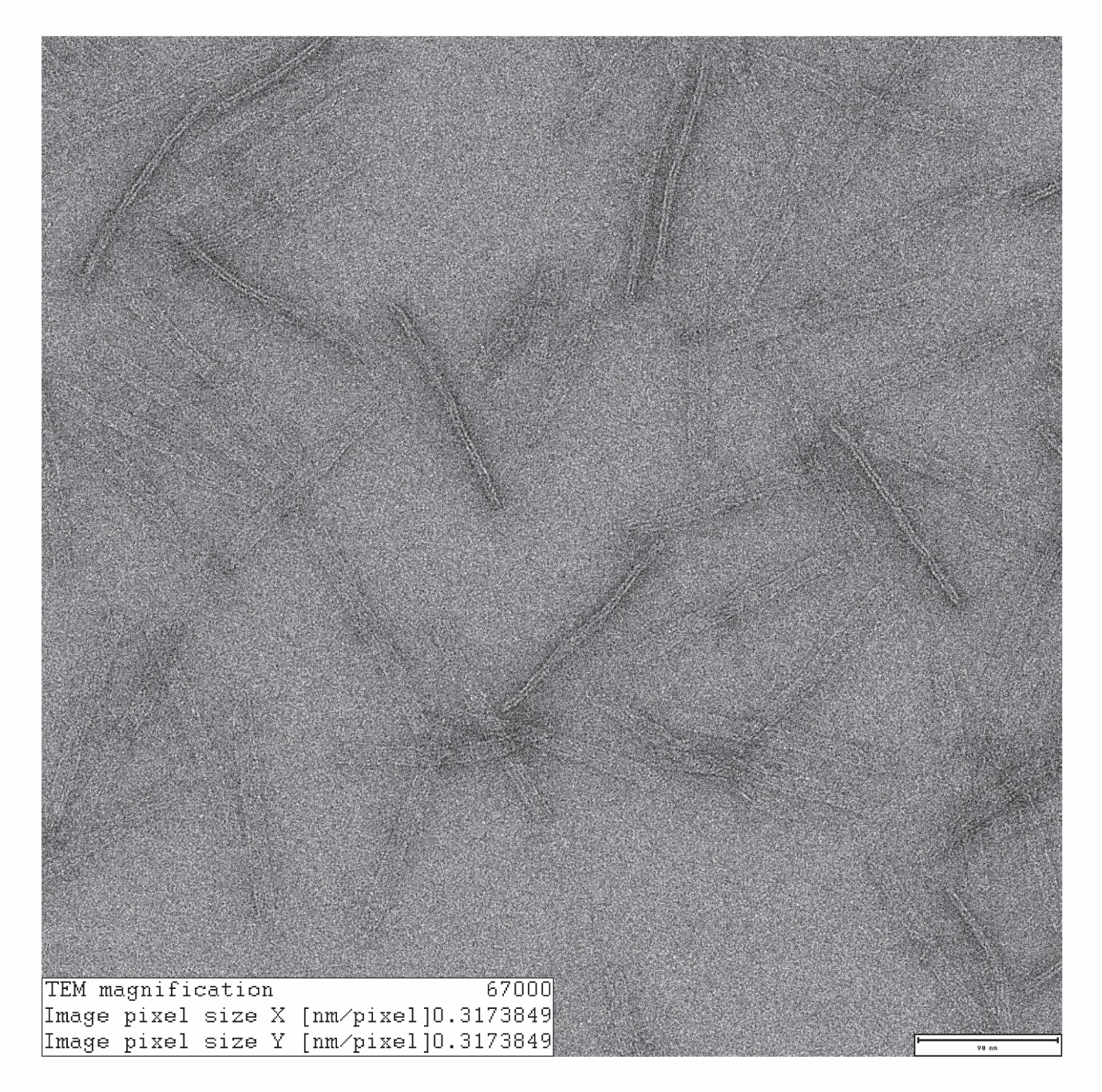

**Supplementary Figure 5.** Full TEM image of the 14HB with hinge 1. Before imaging, the structures were purified from the excess staples via Amicon filtration in a buffer containing 12.5 mM MgCl_2_, 50 mM NaCl, 1x TAE. Scale bar: 90 nm, magnification: 67,000x. A crop-out of this image is used in Figure 1e.

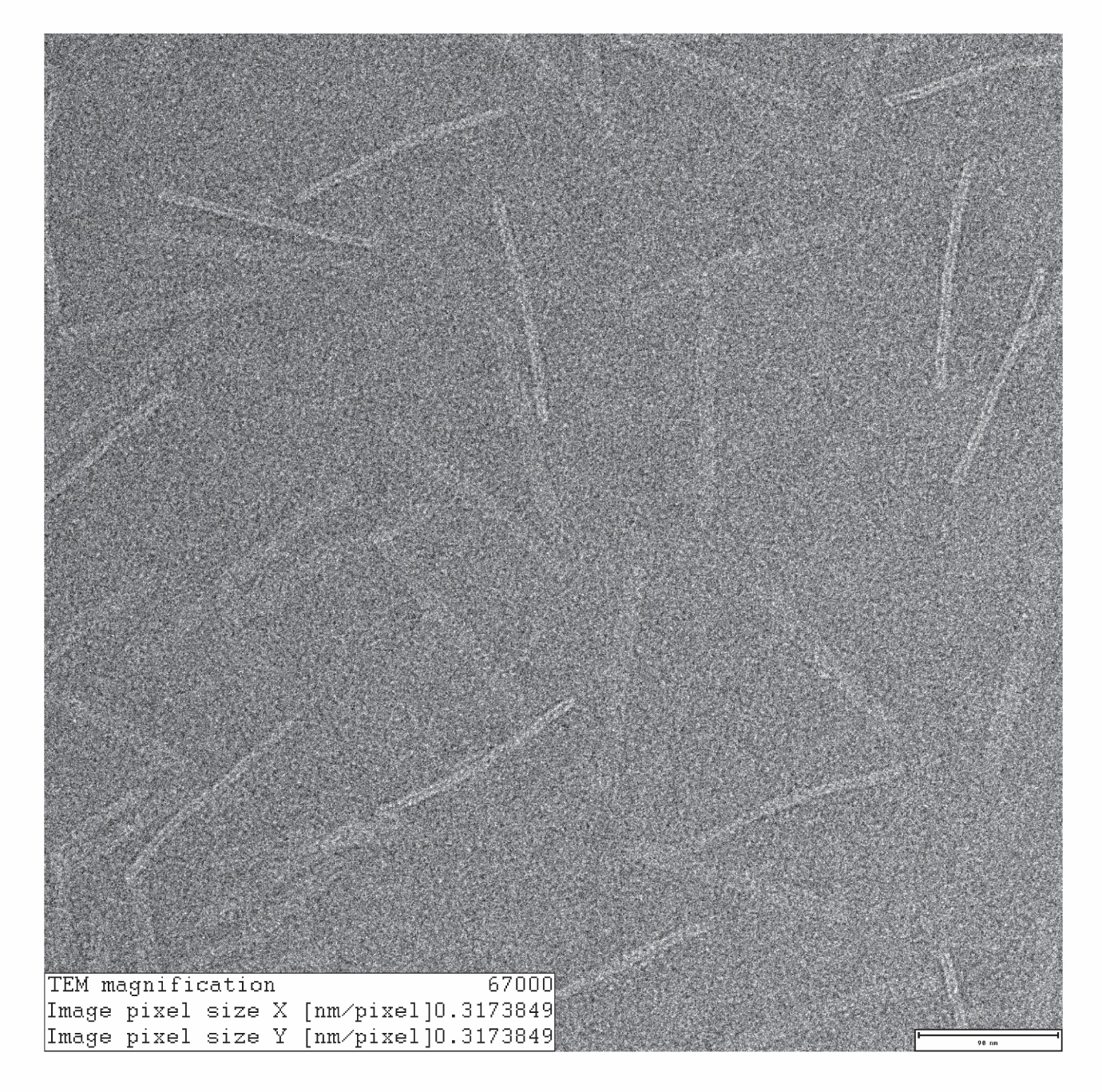

**Supplementary Figure 6.** Full TEM image of the 14HB with hinge 2. Before imaging, the structures were purified from the excess staples via Amicon filtration in a buffer containing 12.5 mM MgCl_2_, 50 mM NaCl, 1x TAE. Scale bar: 90 nm, magnification: 67,000x.

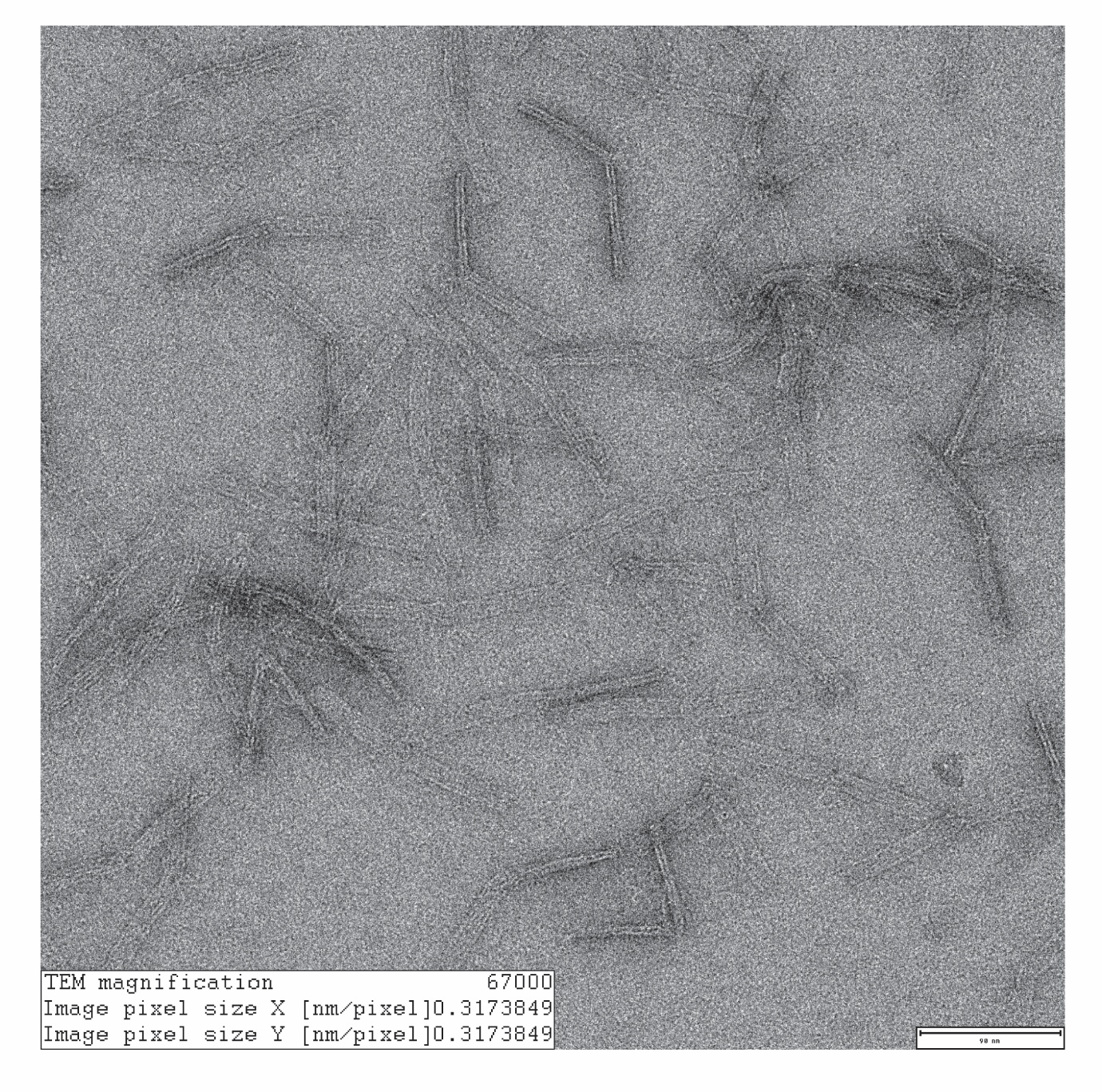

**Supplementary Figure 7.** Full TEM image of the 14HB with hinge 3. Before imaging, the structures were purified from the excess staples via Amicon filtration in a buffer containing 12.5 mM MgCl_2_, 50 mM NaCl, 1x TAE. Scale bar: 90 nm, magnification: 67,000x. A crop-out of this image is used in Figure 1h.

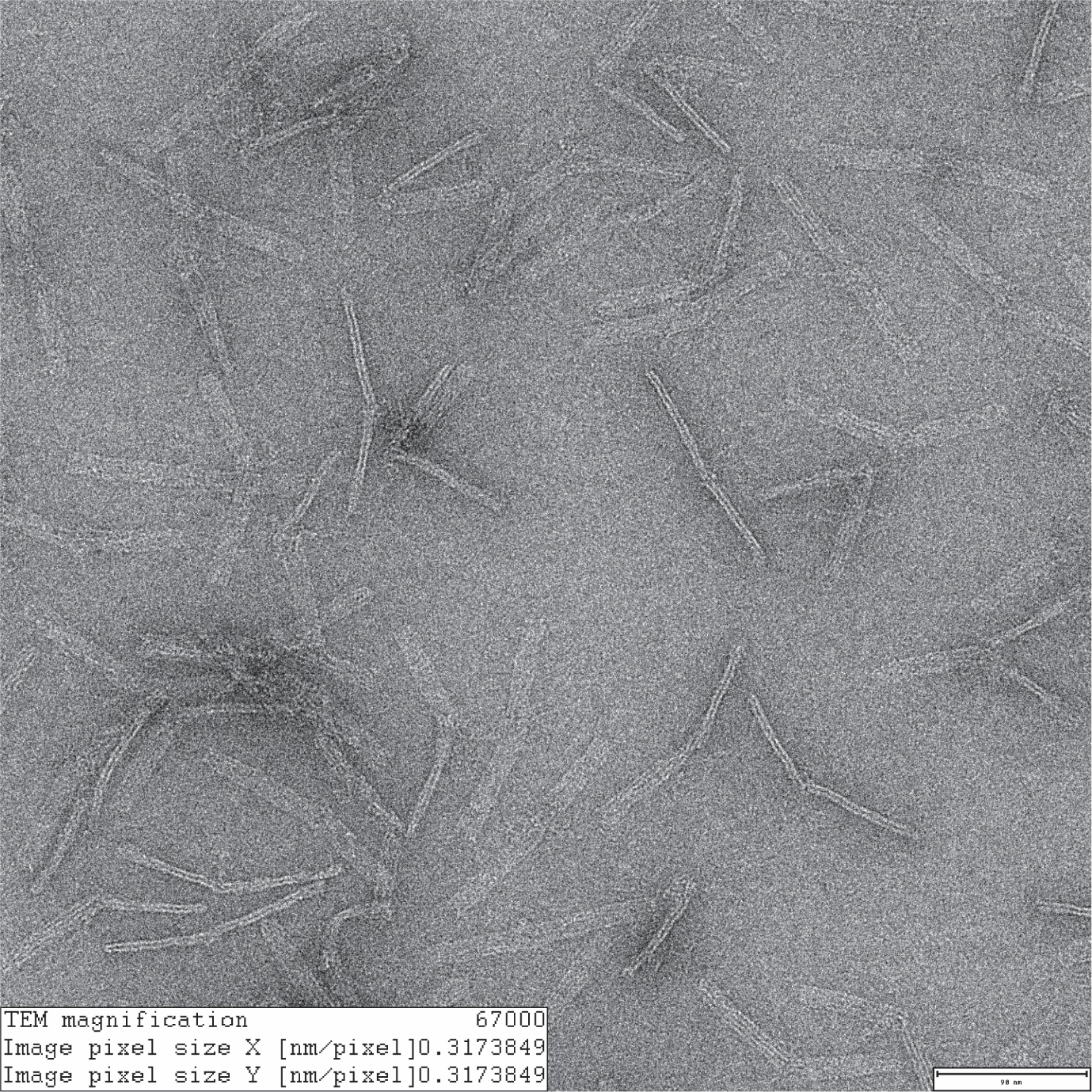

**Supplementary Figure 8.** Full TEM image of the 14HB with hinge 4. Before imaging, the structures were purified from the excess staples via Amicon filtration in a buffer containing 12.5 mM MgCl_2_, 50 mM NaCl, 1x TAE. Scale bar: 90 nm, magnification: 67,000x.

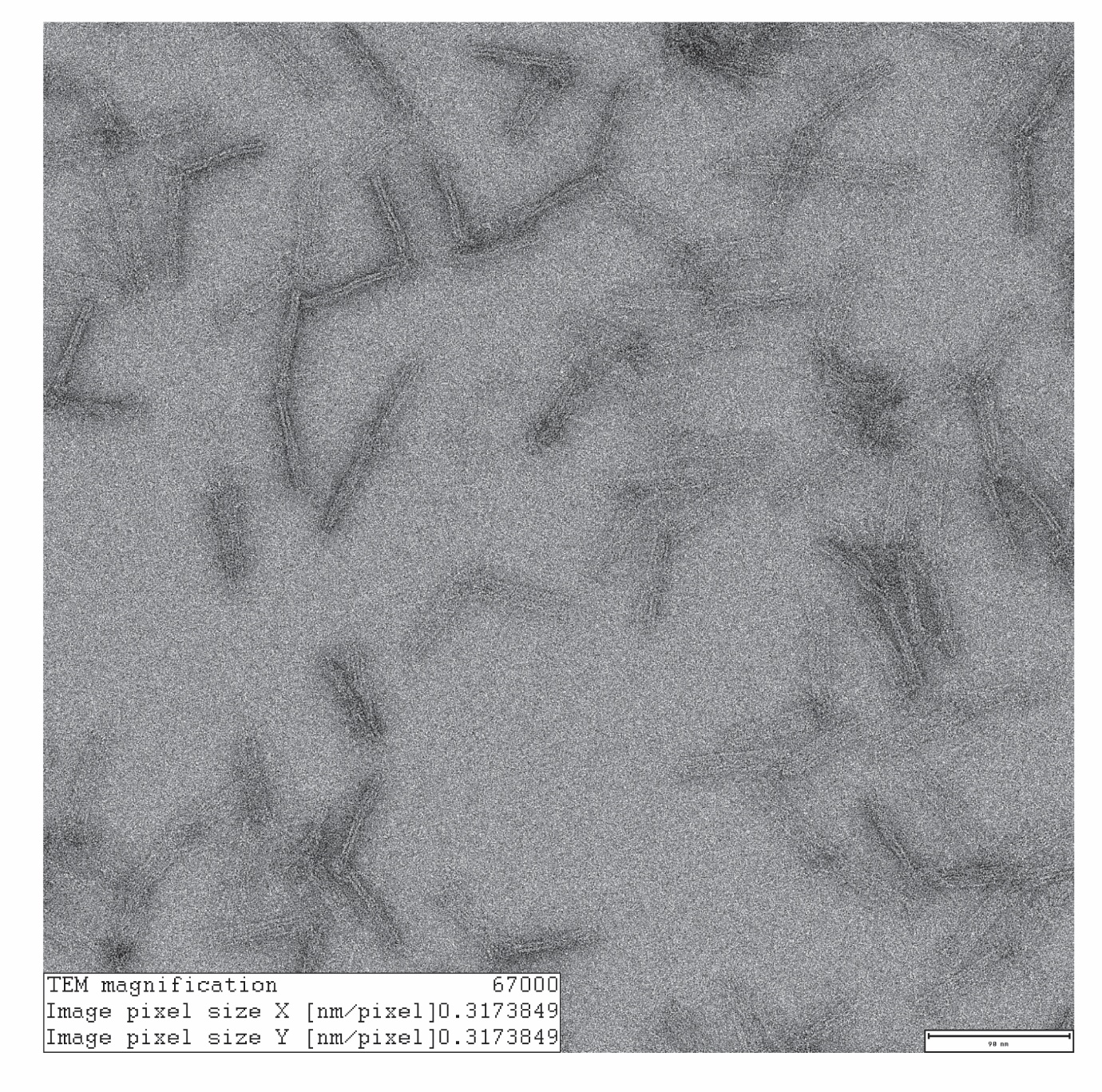

**Supplementary Figure 9.** Molecular weight distribution of proteins in each sample

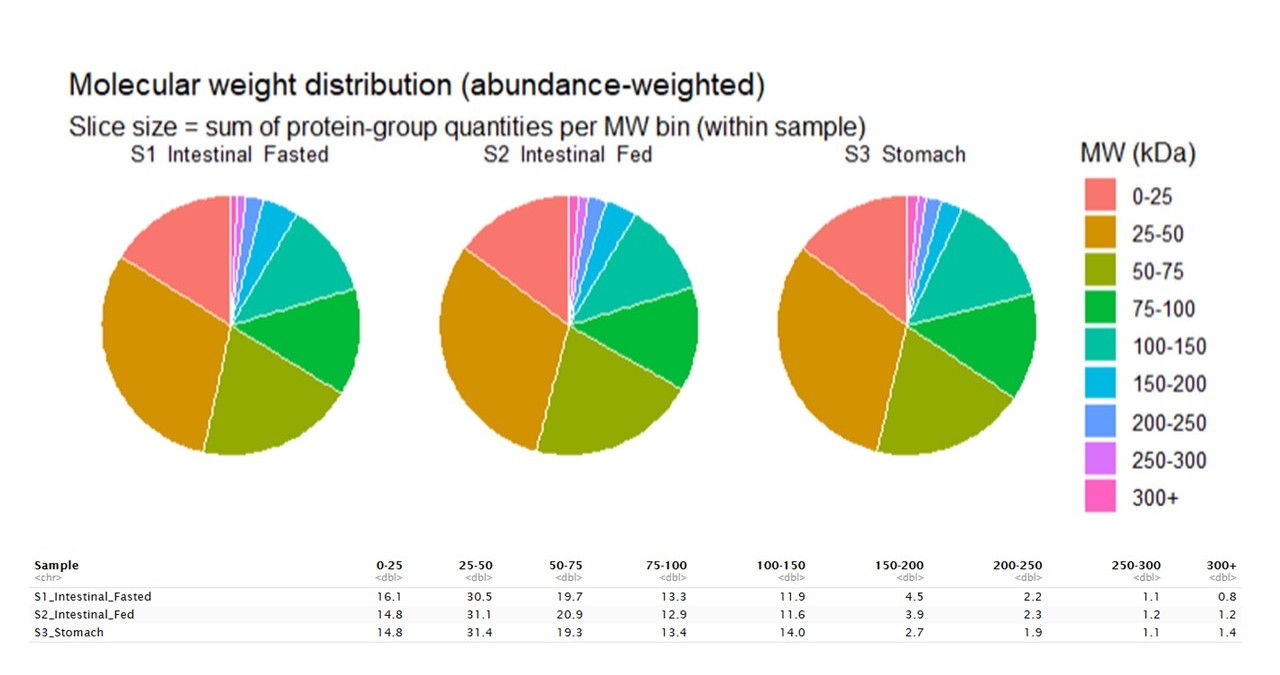

**Supplementary Table 1**. Statistical analysis of the data shown in Figure 2C

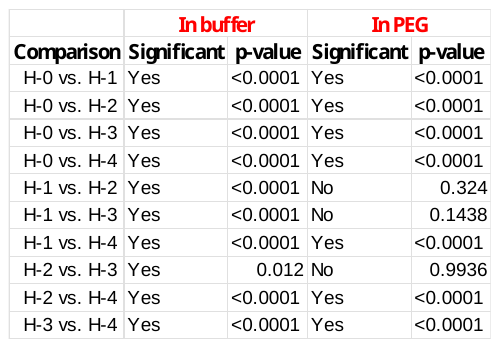

**Supplementary Table 2**. Top 50 proteins contributing to PC1 and PC2.
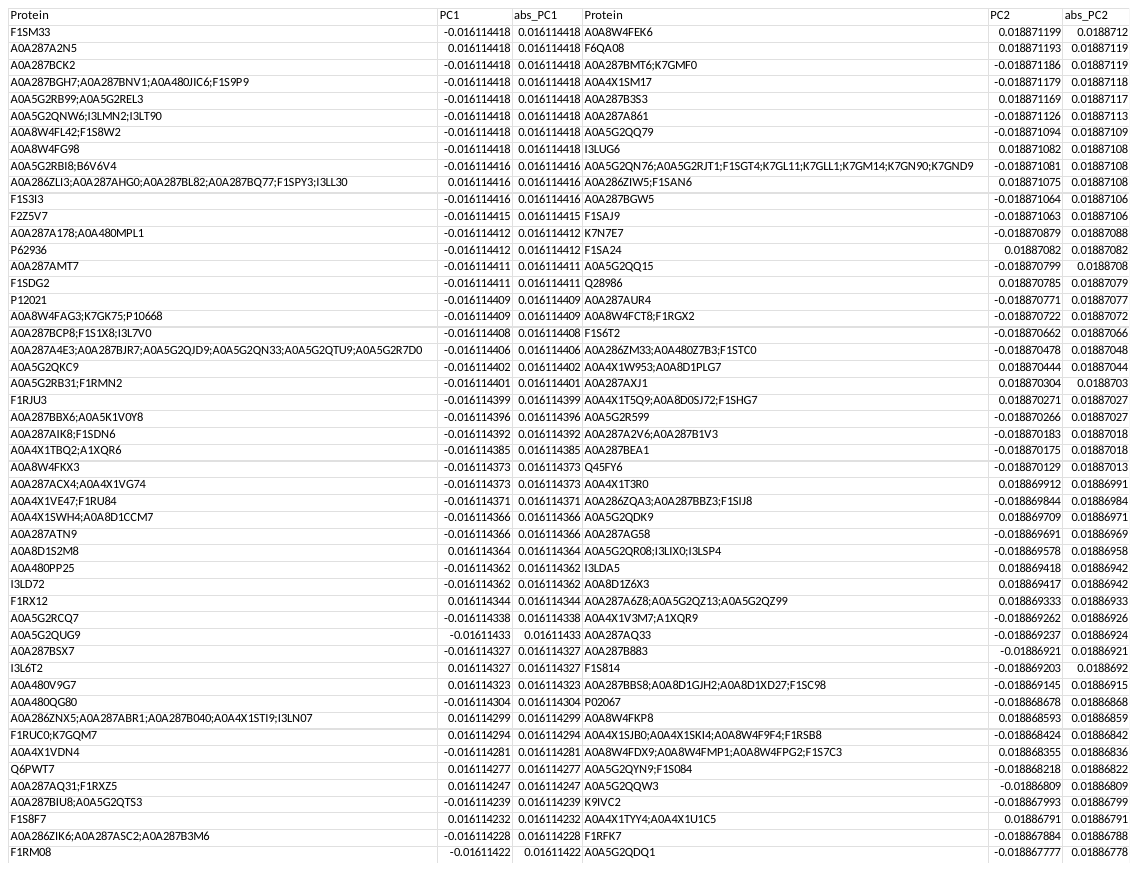

**Supplementary Table 3**. Table of core sequences for constructing the 14HB DNA origami

| Start | Sequence |
| --- | --- |
| 0[83] | CCAGTACCAACTTTGGTTTATCAGCTTGGCCGCTTATCCCTTTTATTTC |
| 0[104] | CCGTAACGAGTGAGAAAGGAGCCTTTAACCCTCAGGGTGGTTAATTTTT |
| 0[146] | CCCTCATTTTCAGGGTAAAGCACTAAATGCAAGCG |
| 0[167] | CCGCCACTTGCCTTAAACCATCGATAGCAAAGACTCTGAGAGAAGGGTG |
| 0[335] | TTAAGAGCCCTCAGACAAAAGGGCGACACGCCTGACATGGAATTTGACG |
| 0[398] | CCCGTATAGGTCAGCCACGGAATAAGTT |
| 0[440] | TGGTAATTCATTAAACATACATAAAGGTCGCAATAATTAGTAGACCTGA |
| 0[461] | TTTGATGGCGCAGTATGTTAGCAAACGTAAAAGAACGTTGTAGTGGCAC |
| 1[28] | GTCTTTCTAGCGTATAAAGAACGTGGACGTTGAGTTATATTCATTATTA |
| 1[71] | GCTAAAAAACTACAACGCCTGTAGCATTCTGTATGGTGAAT |
| 1[133] | GAATAATAAAGTAGTAATCTTTTTTCACGCTACAGGTCCACGAATGTGT |
| 2[34] | ATAGTTGTAACCGAGTTGTTCCAAAAAC |
| 2[153] | AGCACCGCGACAGAATCAAGTCCTCAGACTAAAGG |
| 2[307] | GGAGGGATACAACGGCTTTCCATTAATT |
| 2[370] | CAATAGAACTTAGCCAGAACAATTCACC |
| 2[384] | TATTTTGCAGGAGGTTGAGGCAAACAGTAACAGGA |
| 2[447] | AGAAAATAGCCAGAATGGAAAATACAGGTTATAAT |
| 3[140] | AGGCTTTTAATCATTGTGAATCGAGAGGCTTTTGCTCAAAAA |
| 3[294] | AACAAAGAGGTAAACGCCACCCTCAGAAGATTAGGATGCGCC |
| 3[357] | CCATGTTAAATTCACAGAGCCGCCGCCATATTTCGTTAGAAT |
| 3[420] | GAGGAAAGGCAACAAAACAAATAAATCCAAGTTTTCCAGAAT |
| 4[34] | CAGGTAGCATTCAAGCCTCAGTAAAGTA |
| 4[90] | GACGTTGATCGTCATTGTATCCAACAGTTTCAGCGACTGAGTGTGAACC |
| 4[97] | CAGTCAGTCATAACGTAGCATAGATTTAGTTTGACAAATCCATCATCTT |
| 4[132] | TACCTTAAGCAACGGTTGAAAGAATTGC |
| 4[307] | ATAGGCTCATATTAACCTCAAATCAATA |
| 4[321] | TACAGACGTATCATTTCAACCCCTCAGAGCCACCAGCTGAGAACTATGG |
| 4[328] | AACGGTGAATAAGATCTAAAGTGGATTA |
| 4[370] | CATAAGGCAGCCTTCCTGCAAATCAAAA |
| 4[391] | TAAGAAAAGCAGACAGGCGTAAGCAGATATCGGCCTAATAAA |
| 4[433] | GAAACAACACCCTGAAAATACAGATGAATATACAGCGCAGAGTAGATAA |
| 4[454] | TAAGCCCGTAATTGAGCCCTAACCTTTT |
| 5[42] | AGATACAGAACAACGGTCGCTACAGCTT |
| 5[336] | TTTTGTTAAAGAGGTGTCGAAGCCAAAG |
| 5[385] | CATAAAAAGGCGGTAACAGAAATAAAGAGATGATGCAACATG |
| 5[462] | ATATCAGCACAAGAGATTAAGACGCAGTCTCTGAA |
| 6[55] | ATCGCGTGCTCCTTAGCTTAATTGCTGATGGAAGTAATTAAG |
| 6[69] | GAAAGACATTACGACGTTAATGTTAAAGCTTTCGAGGGATTTT |
| 6[97] | TTGCATCGTTAATTATCGCAAGACAAAGAACGAGT |
| 6[139] | TCAGGTCCGACCGTGGTTGGGTTATATAGGTCAATGAGCTGA |
| 6[160] | AGAATGAGTTAAATACCTTTTTAACCTC |
| 6[307] | TTATCCGAATTTAGATATATGTGAGTGAGCAATTCATATCAA |
| 6[328] | AGCAAATGAGCCAGATGGAAACAGTACAGATTGTTCATCACC |
| 6[391] | TCATCGACAATAAAAAACAAACATCAAGCACGTAACAGTATTCGACCAG |
| 6[433] | TCCTTATTCAACAAGCGAATTATTCATTTAACGTCCGAACGA |
| 7[21] | AGGTCAGAACCAGAAATACCAAAAGATTCCACGCACGCCGAC |
| 7[105] | CTGACCTAGAAGCATACCAGACTGGCTCAGACAGCGGCTCCA |
| 7[168] | AACACCGCAGTTCAGGTAATATGGGCTTGAGGAAGACCAATG |
| 7[273] | ATCGCCATTTTAGCCTAATTTATCTTGACAGCGATATTCATT |
| 7[399] | TTCAGCTACCAAGTAGCGCATACCGAAGCAAAGTTAAAGACA |
| 7[441] | GTCCTGATAATCGGTCAGAGGAATAATAAATACCC |
| 8[27] | CTCAACAATGCAACAGCATAAGTTGTACCAGTTTGCCACTAT |
| 8[111] | CTGATGCCATTAGATTCTACTTCATATAAAATCCTAAATCAACCATGTA |
| 8[146] | CGGCTTAGTGATAAATAAGGCCCATAAAAAAAGAATCAACTTGAGGACT |
| 8[174] | AGGTCTGCTCGTATCATTTGACCGGAGATTCACCGCGATTTACTCAGAA |
| 8[279] | GTAAATCATTCCTGTCTGGTCGGGGTGCAGCTGCACGCGTAAACCAGGC |
| 8[342] | AATTACCGAATAATCAGCAAAGTCTGAAACAGGAAGACGAGCGAAAGTA |
| 8[363] | TACATTTCTACCATCAGTGCCTTGGCAGATATTAC |
| 8[405] | AAAAGAAAATTGCGATAAAACTTCTGGCGAAGAACTAAAGGGAACAGTG |
| 8[447] | AAAATCGTAACAGTAAACATCAGAATACGCAATACAGTGTTTAGTGTAC |
| 8[468] | TTGCTTTGAGAAACATGCGCGATTTTTGTCACGCAGCCACCGCATGGCT |
| 9[35] | CGGTGTCATATAATGAGTACCTTTAATTTTTAATT |
| 9[56] | CATATAAGGCTTAGTTGATAAGAGGTCATTCTTTTGAAAATTTGCGGAT |
| 9[63] | CAGTTGAGGCAAGGGAAGCCTATAAATCACCGTCTATCAGGGCGATGGC |
| 9[77] | TTCTGCGAACGCGATCAAATATATTTTAAAAAAGAAACACTA |
| 9[119] | CGCAAATACTATATATGGTTTGAAATACTTTACCCAAAATAG |
| 9[287] | GATGATGATAACCTCAACGCCAACATGTGTATTCTAAACAGCGGCTGAC |
| 9[308] | TAATCCTTAAATCAGCAGAGGCATTTTCCAGATATCAATCCA |
| 9[329] | TACTTCTTTTTTTATAATAAGAGAATATCGCGCCC |
| 9[350] | TTAGAACAACAATTCGACAAAAGGTAAATTTCATCAAAATAGGAACCGA |
| 9[371] | TTATTTGAAAACAATGTCCAGACGACGAGAACAAGAGAATAA |
| 9[413] | TCAGGTTTCAATTAAACGCGCCTGTTTACATTCCAAACTGAATGAAATA |
| 9[455] | ACATCGGGAATACCAAATAATATCCCATGAGCATG |
| 10[90] | TAACATCGGATAAACCGAAATCCACTACTTCGTCA |
| 10[139] | GGGGCGCAACCTGTTTAGCTATATTTACTAGACTTTTCATTT |
| 10[160] | GAAGTATAAACAATTCGACAAAGAGACTAAGAATA |
| 10[314] | TTGCTGATTTATCCAGAAGGC |
| 11[35] | ATTATGACAATAAACTAATGCCGAGCTTCAAAGCGGATTAGAGCTGTAG |
| 11[42] | CCCTGTAAGATAGGTCCAACGACAGCCCTCATAGTCAGACGTGATACCG |
| 11[56] | TGCGGGACAAAGAAAAAAGGATTCAAAT |
| 11[77] | AACGCAACAATAAATAAGAGCTTAAGAGGAAGCCC |
| 11[98] | AGAACCCAATAGTACCTCGTTAAGCGGA |
| 11[119] | TGCAATGAAAGGTGAAAAACCTGACTATTATAGTCAAATTTAGTAAATG |
| 11[126] | CCTGAGTCTGGTTTAGGTGCCGATAGCAAGCCCAAACTAAAGATCTCCA |
| 11[140] | AGGTAAAGATTCAAAGTTGCACGGAACCGCCACCA |
| 11[161] | AGAAAGGGGATTTACAGAGGGGAAAACG |
| 11[266] | AAAGCCTAGTTGGCCCAGAGCGAACCTCCCGACTTGGGCTTATTAATTA |
| 11[287] | GTGAGCTACCCTCAACAAAATAAGAACG CGAGGCGTATTTAATGCTTCT |
| 11[308] | GCGTTGCGCTACATATACCTCACTGCCCGAGATTTCAGGCGC |
| 11[329] | CTCAATCTGAAAAAAACGATTAATAGCA |
| 11[350] | ATTTACAACGCTGAAAAAATGGTAGGAATCATTACAAAGTACTCATTTG |
| 11[371] | AGTCACAAACACCGTACAGAGCAAGCCGTTTTTATGTAATTCAATTAAT |
| 11[392] | AGGGACAAGAGGTGACAGGGAACCGCAC |
| 11[413] | GATAGAAACCACCAGAGAATTAGAACGGGTATTAAAATGCAGCCTGAGC |
| 11[434] | AAGCGTAGCCATTAAACAAAGCTGTCTT |
| 11[455] | AGACAATAACTGATAGCGCTATAGAAACCAATCAAACAAGAAAAGTTAC |
| 12[83] | CGGCAAATTGCGGGGGAAGAAAAATCTAGGCATAGTCATACATTCCCAA |
| 12[384] | TTGCTGGGGAGCTATAATGCCCCCTGCCGCATTGATCACAAT |
| 13[56] | GCGAAAAAAAAGAAGCAGGGAAAAACGA |
| 13[98] | ATCACCCGTTTGATCAGCGAAATTATAC |
| 13[119] | GGGGTCGGCCCCAGCGAGGGTTGCGATT |
| 13[161] | GAGCCCCCCTGGCCTTTTCATGAGATGGTTTAATTGTTTTGC |
| 13[182] | ACGGGGAACAGCTGTAAACGGAACGAGT |
| 13[266] | CACGCTGTTAATGATGACCCCCAAGAAC |
| 13[287] | ACCCGCCAAACCTGAGCGCGACTTCATC |
| 13[308] | GCTACAGGGCGCGTCTCCTCAAGAGAAGCCGCCACGATTGAG |
| 13[329] | TTGCTTTAAACGCTTAAATTGACAGATG |
| 13[350] | ACGTGCTCGCCAGCACCTGCTACTGACC |
| 13[357] | TTCCTCGGAACCTATTATTCTCCACCACTATGGTT |
| 13[371] | CAGAGCGTAATATCCGGAACGGGTCAAT |
| 13[392] | GGCCGATTCAAACTAGCCGAACCCTTTT |
| 13[413] | ACAGGAACACTTGCGGAAACCGCAATAG |
| 13[434] | CCTGAGATTCTTTGATAACGGAGAGCAA |
| 13[455] | CAGTGAGAATTAACCTGGCATATTGAGT |
| 1[168] | TAGCGTCAGACTGTCGCCACCGAGCTTG |
| 1[273] | CGGAACCGCCTCCCGCTCAGTCCACCAC |
| 1[336] | AGCCGCCACCAGAAGAAACATACGTATA |
| 1[399] | ACGATTGGCCTTGACTTGAGTATTTTAG |
| 2[55] | TTCTTAAGAGGCTTTAGCCCGATACTTT |
| 2[118] | AAAAAAAATCGGAACAGGCGATTTTAAA |
| 2[181] | AAACGTCTTTCCATATTGCCCCAGTCAA |
| 2[286] | GGAAATTTATACCATCGTGCCCTAATGA |
| 2[349] | TACCAGCATCCGCGCATTGCAATGGATT |
| 2[412] | GAAACGCACCAGAACTGAGTACAACAGA |
| 11[182] | CTTTAAAGAATCATAAATCAT |
| 11[189] | CGAGGCGTATTTAATGCTTCT |
| 11[287] | GTATTAAAATGCAGCCTGAGC |
| 11[294] | TCAATATACGGGCAAAGCCGG |
| 11[413] | AACTCACAGTCGGGGCGCTTA |
| 11[420] | CCCTTCTATAACATCGGTACG |
| 11[189] | ATCACCAGTCAATAGTTTAGATCAAATG |

**Supplementary Table 4**. Table of staples for generating the hinge.

Hinge level 0 contains all staples. Hinge level 1 excludes hinge level 1 staples. Hinge level 2 excludes hinge level 1 and 2 staples etc. The removal of staples generates a wider hinge, enabling more flexibility.

| Start | Sequence |
| --- | --- |
| Hinge 1 - Staple 1 | TCATTTTGCGATAGAATTCTTACCAGTAAATCAAGTATCCTGCAAAGCT |
| Hinge 1 - Staple 2 | TGCGTATGGAGCGGGCCCGGAATAGGTGGCGTTTGTTAGAGC |
| Hinge 1 - Staple 3 | AGCGAAATGGGCGCTAAAACGGCTCATTCAGTGAA |
| Hinge 2 - Staple 1 | CAGCAAAACCAACCCAGGGTGATAAATTAATGCAATTCCACA |
| Hinge 2 - Staple 2 | AAAGAGGTTGGGAACCATCTTTTCATAAAAGTATAGCGCTAG |
| Hinge 2 - Staple 3 | TAAGGCTCGAAGGCATCACCATAGCCCCCTTATTATATCACCGGAAGAA |
| Hinge 2 - Staple 4 | AACGTAAAATCTTATGAGGAAAAAGAAA |
| Hinge 2 - Staple 5 | GCTATTTTGCGTCAGGAATCACCCAGCTTCTTTAGAACATTA |
| Hinge 2 - Staple 6 | AAACATAGCGGAACGGTTATCTAAAATAACAATTTATTAGTT |
| Hinge 2 - Staple 7 | GAGCCATCAAAAGAGGCGGTTCAACATA |
| Hinge 3 - Staple 1 | GGAGGTTTCGGTCAGTAGCACCATTACCACGTAATCAGTGAGGATATTC |
| Hinge 3 - Staple 2 | GAGAGGGTTGATATTCAAAATCCGACTT |
| Hinge 3 - Staple 3 | TAAGACGTTTGAGTGAGCACTCTAGCTGGTTTTTCAAGGAAGGTACTCA |
| Hinge 3 - Staple 4 | TTAAAAGCTGAGAATAGTATCATATGCGTTATACACTTAGAT |
| Hinge 3 - Staple 5 | CCACCAGCTTAGAAAACGCTCAACAGTAGCGGGAGTAACGAG |
| Hinge 3 - Staple 6 | AACCGTTAACAACTATACTGCTAAATAT |
| Hinge 3 - Staple 7 | CGAGCCGAAGGAATCCAACGCGTTTTGAAGCCTTATAAAGCCTCCTTGA |
| Hinge 3 - Staple 8 | GGCGAGATTTTCACGCCACTATGCCCTG |
| Hinge 3 - Staple 9 | GGCGCTGGGGGAGAATACACTCCAAATC |
| Hinge 4 - Staple 1 | GGATAAGCCACCACAAAGGTGAATTATCTCATCTTATCGGCCAAAGTGT |
| Hinge 4 - Staple 2 | ACACCAGGTAAAATATTAGCATTTCATCGGCATTTTAGTACCCGAACGT |
| Hinge 4 - Staple 3 | ACGAGAAGCGTCCAAATAGATATTAATT |
| Hinge 4 - Staple 4 | TCATTACAAAACACACCGTCACACCGGAACCAGAGTGCCGTCTAGCGGT |
| Hinge 4 - Staple 5 | CGGATATCGTCTTTAAATCAAGGAATTATCATCATGTCGCTAATTGAGA |
| Hinge 4 - Staple 6 | TCATTGAGCCTGTTGAGTCAATAGTGAAGAACGTTTAGAGCC |
| Hinge 4 - Staple 7 | ATTTTCCAAGGAGCCAGTTGAGAAGCATAACGCGCGCAAGTG |
| Hinge 4 - Staple 8 | TTTGCCCTTTATCAAATTACTAGAAAAAATCCCCCCTGGATA |

**Supplementary Table 5.** Table of staple sequences for the attachment of ATTO647N to the DNA origami.

For the single molecule tracking experiments, 6 staples with an extension for capturing the ATTO647N-functionalized DNA strand was used. In this case, staples 1-6 with the extension (in red) were used for annealing the structure. If no ATTO fluorophores need to be captured, the unfunctionalized staples (UF) with no capture extension were used.

| **Start** | **Sequence** |
| --- | --- |
| UF ATTO 1 | ACTAACGTAACGCCTTAGCAATTCATTC |
| UF ATTO 2 | TTAAGAACGACGATGCATCAATACATTT |
| UF ATTO 3 | AGTAAATGTAAAATGATAATATAAATCC |
| UF ATTO 4 | AAGAGTAGCCAGTTATCAATAATTATCA |
| UF ATTO 5 | AACTTTGTAACGTCGAGCCAGGGAAGGG |
| UF ATTO 6 | CTATCTTTAGACGGGCAGAAGTAGATTT |
| ATTO 1 | ACTAACGTAACGCCTTAGCAATTCATTCGGGCTCATGCGAGGCTGTATGT |
| ATTO 2 | TTAAGAACGACGATGCATCAATACATTTGGGCTCATGCGAGGCTGTATGT |
| ATTO 3 | AGTAAATGTAAAATGATAATATAAATCCGGGCTCATGCGAGGCTGTATGT |
| ATTO 4 | AAGAGTAGCCAGTTATCAATAATTATCAGGGCTCATGCGAGGCTGTATGT |
| ATTO 5 | AACTTTGTAACGTCGAGCCAGGGAAGGGGGGCTCATGCGAGGCTGTATGT |
| ATTO 6 | CTATCTTTAGACGGGCAGAAGTAGATTTGGGCTCATGCGAGGCTGTATGT |
| ATTO647N strand | ACATACAGCCTCGCATGAGCCC-/3ATTO647NN/ |

**Supplementary Table 6.** Table of staple sequences for the attachment of protein-DNA conjugates to the DNA origami.

When no enzymes are needed on the origami, all 4 unfunctionalized (UF) staples were used 4. For structures with four capture strands, staples 1-4 with capture extensions were used.

| Start | Sequence |
| --- | --- |
| UF Staple 1 | TAGTAAATGAATTTTCCACAGTCAAAGG |
| UF Staple 2 | AATAGAAAGGAACATAGGAACGTTTTTT |
| UF Staple 3 | ATTAGCGGGGTTTTTCAGAGCTATTGAC |
| UF Staple 4 | AACGGGGTCAGTGCTATTCACTATAAAA |
| F Staple 1 | TAGTAAATGAATTTTCCACAGTCAAAGGAAAATTCAGGATTCTCAATT |
| F Staple 2 | AATAGAAAGGAACATAGGAACGTTTTTTAAAATTCAGGATTCTCAATT |
| F Staple 3 | ATTAGCGGGGTTTTTCAGAGCTATTGACAAAATTCAGGATTCTCAATT |
| F Staple 3 | AACGGGGTCAGTGCTATTCACTATAAAAAAAATTCAGGATTCTCAATT |
| Protein strand - NHS | /5AmMC6/ AATTGAGAATCCTGAATTTT |
